# Presynaptic PTPσ organizes neurotransmitter release machinery at excitatory synapses

**DOI:** 10.1101/2020.01.10.901546

**Authors:** Kyung Ah Han, Hee-Yoon Lee, Dongseok Lim, Jungsu Shin, Taek Han Yoon, Chooungku Lee, Jeong-Seop Rhee, Xinran Liu, Ji Won Um, Se-Young Choi, Jaewon Ko

## Abstract

Leukocyte common antigen-related receptor tyrosine phosphatases (LAR-RPTPs) are evolutionarily conserved presynaptic organizers. The synaptic role of vertebrate LAR-RPTPs *in vivo*, however, remains unclear. This study systematically analyzed the effects of genetic deletions of LAR-RPTP genes by generating single conditional knockout (cKO) mice targeting PTPσ and PTPδ. Although the numbers of synapses were reduced in cultured neurons deficient in individual PTPs, abnormalities in synaptic transmission, synaptic ultrastructures, and vesicle localization were observed only in PTPσ-deficient neurons. Strikingly, loss of presynaptic PTPσ reduced neurotransmitter release prominently at excitatory synapses, concomitant with drastic reductions in excitatory innervations onto postsynaptic target areas *in vivo*. However, postsynaptic PTPσ deletion had no effect on excitatory synaptic strength. Furthermore, conditional deletion of PTPσ in ventral CA1 specifically altered anxiety-like behaviors. Taken together, these results demonstrate that PTPσ is a *bona fide* presynaptic adhesion molecule that controls neurotransmitter release and excitatory inputs.

## Introduction

Distinct molecular assemblies at presynaptic nerve terminals and postsynaptic densities are responsible for the fast and precise transmission of neural information (Südhof, 2018). These structures act by coordinating the regulation of bidirectional signals across synaptic clefts, determining the properties of individual synapses, including the computation of neural information (Südhof, 2018). Several synaptic cell-adhesion molecules are thought to act not only as physical connectors across synaptic clefts, but also as *trans*-synaptic signaling hubs (Missler *et al*, 2012, Südhof, 2017, Südhof, 2018). Neurexins (Nrxns) are evolutionarily conserved receptors, with extensive splicing at six canonical sites (Aoto *et al*, 2015, Aoto *et al*, 2013, Dai *et al*, 2019). The splicings regulate the binding of Nrxns to distinct ligands, as well as glutamate receptor-mediated postsynaptic responses and long-term synaptic plasticity (Aoto *et al*, 2015, Aoto *et al*, 2013, Dai *et al*, 2019). Conditional knockout (cKO) mice with complete ablation of all *Nrxn* genes showed extreme phenotypes, depending on synapse types and analyzed brain areas (Chen *et al*, 2017). These findings suggested that Nrxns are context-specific regulators of synapse properties, but are not essential for synapse initiation (Chen *et al*, 2017).

Similar to Nrxns, leukocyte common antigen-related receptor tyrosine phosphatases (LAR-RPTPs) are evolutionarily conserved key synaptic organizers expressed in presynaptic active zones (AZs) (Han *et al*, 2019, Südhof, 2012, Um & Ko, 2013). Invertebrate LAR-RPTP orthologs (dLAR in *Drosophila melanogaster* and PTP-3 in *Caenorhabditis elegans*) were shown to be expressed in axons/growth cones, playing critical roles in axon guidance, dendritic growth, and synapse formation (Ackley *et a*l, 2005, Chagnon *et al*, 2004, Han *et al*, 2019). In contrast, vertebrate LAR-RPTPs, consisting three members (LAR, PTPσ, and PTPδ), are present in both dendritic spines and axons of cultured neurons (Han *et al*, 2018, Takahashi *et al*, 2012, Wyszynski *et al*, 2002). Analogous to Nrxns, LAR-RPTPs bind postsynaptic ligands, which do not overlap with Nrxn ligands, to induce presynaptic differentiation (Choi *et al*, 2016, Han *et al*, 2018, Li *et al*, 2015, Takahashi *et al*, 2011, Yim *et al*, 2013, Yoshida *et al*, 2011). Constitutive KO mice of individual or multiple LAR-RPTP exhibit pleiotropic abnormalities in both the peripheral and central nervous systems, and impairment in certain aspects of synapse development and function (Elchebly *et al*, 1999, Horn *et al*, 2012, McLean *et al*, 2002, Thompson *et al*, 2003, Uetani *et al*, 2006, Uetani *et al*, 2000, Wallace *et al*, 1999). In contrast, loss of PTPσ and/or PTPδ, using short-hairpin-mediated knockdown (KD)-mediated manipulations, impairs structural and functional development, as well as the LAR-RPTP ligand-induced formation of artificial synapses (Yim *et al*, 2013; Han *et al*, 2018; Takahashi *et al*, 2012; Dunah *et al*, 2005). Intriguingly, PTPσ and PTPδ (but not LAR) serve as functional receptors for presynaptic assembly at specific synapse types (i.e., PTPσ for excitatory synapses, such as Slitrks, TrkC, and SALMs, and PTPδ for inhibitory synapses, such as Slitrk3, and excitatory synapses, such as IL1RAPL1) (Choi *et al*, 2016, Han *et al*, 2018, Li *et al*, 2015, Takahashi *et al*, 2012, Valnegri *et a*l, 2011, Yim *et al*, 2013, Yoshida *et al*, 2011). Although these findings clearly indicated that LAR-RPTPs may be central components in both pre- and postsynaptic neurons that organize various aspects of synapse development, a sophisticated approach using cKO deficient in LAR-RPTPs is required to precisely assess the synaptic role of vertebrate LAR-RPTPs *in vivo*.

The present study describes the generation of mutant mice carrying cKO alleles of PTPσ or PTPδ. We found that, in keeping with the KD effects (Han *et al*, 2018), conditional genetic deletions of PTPσ and PTPδ reduced the numbers of synapses at excitatory and inhibitory synapses, respectively, with PTPσ-deficient synapses showing concomitant impairment in excitatory synaptic transmission. Moreover, deletion of PTPσ resulted in an abnormal increase in the length of presynaptic AZ and in postsynaptic density (PSD), and abnormal vesicular organization in presynaptic boutons. Furthermore, PTPσ loss from layer II of medial prefrontal cortical (mPFC) neurons and hippocampal CA1 pyramidal neurons selectively impaired innervation and neurotransmitter release at excitatory, but not inhibitory, synapses formed on layer V of mPFCs and subicular pyramidal neurons. Strikingly, loss of PTPσ did not alter postsynaptic AMPA receptor-mediated responses, suggesting that PTPσ primarily serves as a presynaptic adhesion molecule that controls the innervation of presynaptic excitatory inputs and neurotransmitter release *in vivo*. Despite PTPσ playing a similar presynaptic role at mPFCs and hippocampal circuits, PTPσ deletions in ventral CA1 selectively influenced anxiety-like, but not social, behavior. These results suggested that PTPσ is essential for the regulation of presynaptic functions *in vivo*, distinct from the roles of presynaptic Nrxns.

## Results

### Generation of PTPσ and PTPδ cKO mice

The goal of the present study was to determine the *in vivo* synaptic functions of vertebrate LAR-RPTP isoforms by examining the phenotypes resulting from the deletion of individual LAR-RPTPs. Studies in constitutive KO mice have precluded investigations of the mechanisms of action of LAR-RPTPs because of the pleiotropic phenotypes that are unlikely unrelated to their synaptic roles (Um & Ko, 2013). Transgenic mice with deletion of *PTPσ* or *PTPδ* were generated by crossing *PTPσ*^f/f^ mice, with exon 4 flanked by loxP sites, or *PTPδ*^f/f^ mice, with exon 12 flanked by loxP site, with a Cre recombinase driver line under control of the Nestin promoter (Nestin-Cre) (**Fig EV1A** and **EV1B**). *LAR*^f/f^ mice were not generated because of the ubiquitous expression of this gene outside the brain (Kwon *et al*, 2010, Um & Ko, 2013). RNAscope-based fluorescence *in situ* hybridization showed that the expression patterns of mRNAs encoding all three LAR-RPTP family members overlap in both the mPFC and the hippocampus (**Appendix Fig 1A**). Both PTPσ and PTPδ mRNAs were detected in *CaMKIIα*-positive glutamatergic neurons and in *Gad1*-positive GABAergic neurons of adult mouse brains (**Appendix Fig 1B** and **1C**).

**Figure 1.**
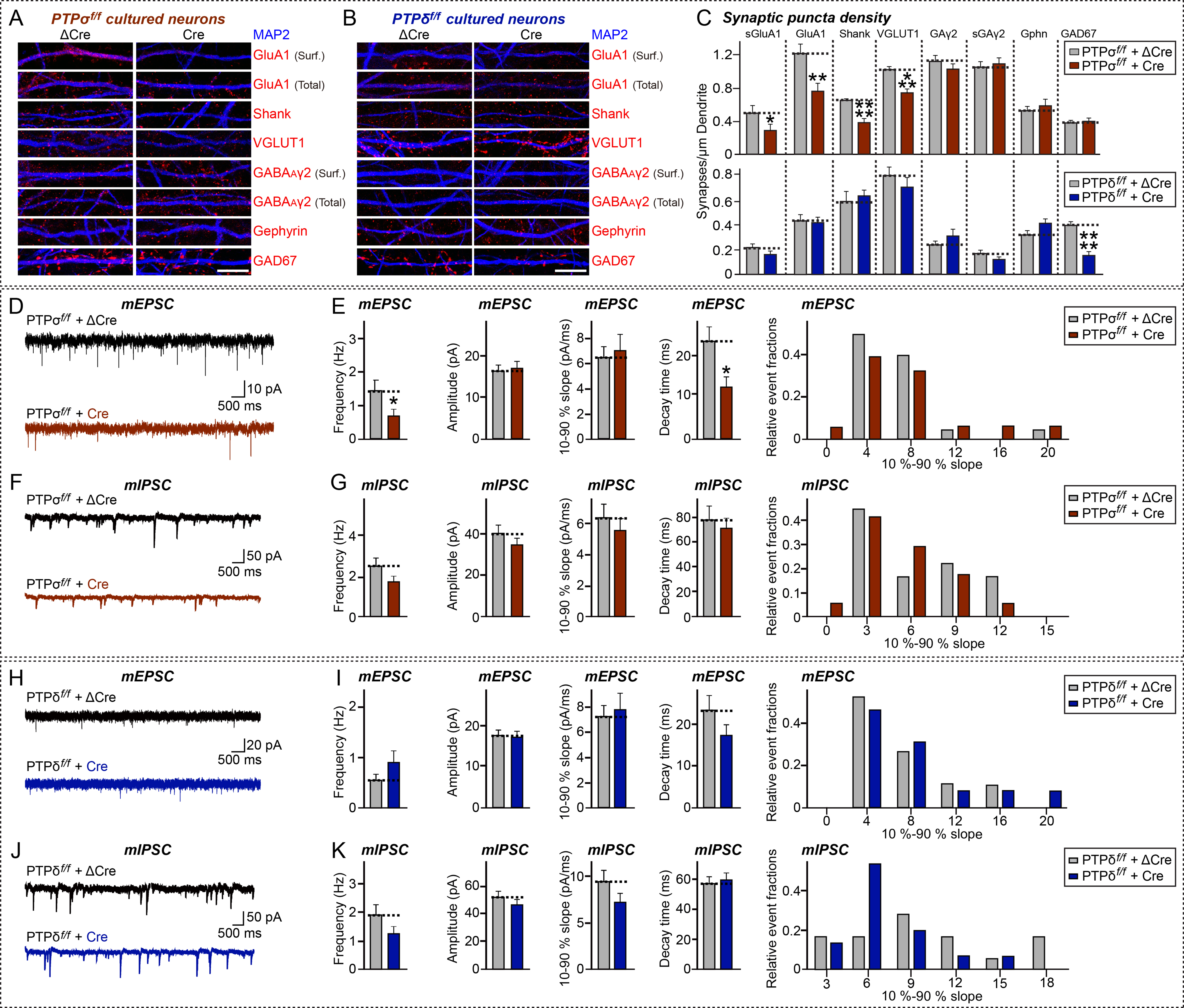
Conditional KO of PTPσ impairs excitatory synapse development and transmission in cultured hippocampal neurons. **A, B** PTPσ cKO (**A**) but not PTPδ cKO (**B**) in cultured hippocampal neurons specifically reduces excitatory synapse density. Double immunofluorescence analysis of MAP2 (blue) and the indicated synaptic markers (red) in mature cultured neurons (DIV14) derived from *PTPσ^f/f^* or *PTPδ^f/f^* mice infected with lentiviruses expressing ΔCre or Cre at DIV3. Synaptic markers assayed included surface GluA1 (sGluA1), total GluA1, Shank, and VGLUT1 as excitatory synaptic markers, and surface GABA_A_γ2 (sGAγ2), total GABA_A_γ2, gephyrin (Gphn), and GAD67 as inhibitory synaptic markers. Scale bar: 10 μm. **C** Quantification of images in (**A** and **B**), measuring the density of the indicated synaptic markers. Data are means ± SEMs (n denotes number of analyzed neurons; ΔCre/PTPσ cKO/sGluA1, n = 16; Cre/PTPσ cKO /sGluA1, n = 17; ΔCre/PTPσ cKO/GluA1, n = 16; Cre /PTPσ cKO/GluA1, n = 15; ΔCre/PTPσ cKO/Shank, n = 16; Cre/PTPσ cKO/Shank, n = 16; ΔCre/PTPσ cKO/VGLUT1, n = 15; Cre/PTPσ cKO/VGLUT1, n = 16; ΔCre/PTPσ cKO/sGABA_A_Rγ2, n = 15; Cre/PTPσ cKO/sGABA_A_Rγ2, n = 15; ΔCre/PTPσ cKO/GABA_A_Rγ2, n = 16; Cre/PTPσ cKO/GABA_A_Rγ2, n = 16; ΔCre/PTPσ cKO/Gephyrin, n = 16; Cre/PTPσ cKO/Gephyrin, n = 15; ΔCre/PTPσ cKO/GAD67, n = 16; Cre/PTPσ cKO/GAD67, n = 16; ΔCre/PTPδ cKO/sGluA1, n = 14; Cre/PTPδ cKO/sGluA1, n = 14; ΔCre/PTPδ cKO/GluA1, n = 14; Cre/PTPδ cKO/GluA1, n = 15; ΔCre/PTPδ cKO/Shank, n = 21; Cre/PTPδ cKO/Shank, n = 21; ΔCre/PTPδ cKO/VGLUT1, n = 15; Cre/PTPδ cKO/VGLUT1, n = 16; ΔCre/PTPδ cKO/sGABA_A_Rγ2, n = 16; Cre/PTPδ cKO/sGABA_A_Rγ2, n = 17; ΔCre/PTPδ cKO/GABA_A_Rγ2, n = 10; Cre/PTPδ cKO/GABA_A_Rγ2, n = 10; ΔCre/PTPδ cKO/Gephyrin, n = 16; Cre/PTPδ cKO/Gephyrin, n = 16; ΔCre/PTPδ cKO/GAD67, n = 16; and Cre/PTPδ cKO/GAD67, n = 16. Mann-Whitney U test; \**p* < 0.05; \*\**p* < 0.01; \*\*\**p* < 0.001; \*\*\*\**p* < 0.0001). **D, E** Representative mEPSC traces (**D**) and quantification of frequencies, amplitudes and kinetics (**E**) of mEPSCs recorded from hippocampal cultured neurons derived from *PTPσ^f/f^* mice infected with lentiviruses expressing inactive (ΔCre) or active (Cre) cre-recombinase. Data are means ± SEMs (n denotes number of analyzed neurons; ΔCre, 20 and Cre, 15; Unpaired t-test; \**p* < 0.05) **F, G** Representative mIPSC traces (**F**) and quantification of frequencies, amplitudes and kinetics (**G**) of mIPSCs recorded from hippocampal cultured neurons derived from *PTPσ^f/f^* mice infected with lentiviruses expressing ΔCre or Cre. Data are means ± SEMs (n denotes number of analyzed neurons; ΔCre, 18 and Cre, 17; Unpaired t-test) **H, I** Representative mEPSC traces (**H**) and quantification of frequencies, amplitudes and kinetics (**I**) of mEPSCs recorded from hippocampal cultured neurons derived from *PTPδ^f/f^* mice infected with lentiviruses expressing ΔCre or Cre. Data are means ± SEMs (n denotes number of analyzed neurons; ΔCre, 19 and Cre, 13; Unpaired t-test) **J, K** Representative mIPSC traces (**J**) and quantification of frequencies, amplitudes and kinetics (**K**) of mIPSCs recorded from hippocampal cultured neurons derived from *PTPδ^f/f^* mice infected with lentiviruses expressing ΔCre or Cre. Data are means ± SEMs (n denotes number of analyzed neurons; ΔCre, 18 and Cre, 16; Unpaired t-test).

To evaluate the cellular effects of endogenous PTPσ and PTPδ deletions, hippocampal neurons were cultured from PTPσ and PTPδ cKO mice. Neurons cultured for at 3–4 days *in vitro* (DIV) were infected with lentiviruses expressing EGFP-fused nuclear Cre-recombinase, which results in a global loss of selective PTP isoforms in all neurons due to high infection efficiency, or with a non-functional mutant version of Cre-recombinase (ΔCre). Global expression of Cre-recombinase caused a nearly complete and specific loss of expression of the target PTP mRNAs in corresponding PTP cKO neurons when analyzed at DIV13–14 (**Fig EV1C**), and specific loss of PTP protein in corresponding cKO brains (**Fig EV1D**). cKO mice in which PTPσ or PTPδ was deleted from the entire brain were viable and fertile, although a modest reduction of body size and birth ratio was observed (**Fig EV1E** and **EV1F**). In addition, PTP-cKO brains showed normal gross morphology, as revealed by staining for the neuron-specific marker NeuN (**Appendix Fig 2A** and **2B**) and for Nissl (**Appendix Fig 2C** and **2D**).

### Conditional PTPσ and PTPδ KO reduces the number of distinct synapse types

To assess the synaptic role of PTPσ and PTPδ, cultured hippocampal PTP-cKO neurons were infected with lentiviruses expressing either ΔCre (Control) or wild-type Cre recombinase at DIV3–4, and the neurons were stained with antibodies to various excitatory and inhibitory synaptic markers at DIV14–16 (**Fig 1A–1C**). The density of excitatory, but not of inhibitory, synaptic puncta was significantly reduced, as measured by staining of PTPσ-deficient neurons with antibodies to GluA1 (both surface and total), pan-Shank, and VGLUT1 (∼30– 40%) (**Fig 1A** and **1C**). Surprisingly, there were no marked changes in the density of excitatory and inhibitory synaptic puncta, except for that of GAD67 (∼50%), on PTPδ KO neurons (**Fig 1B** and **1C**). Moreover, measurements of the apparent sizes of synaptic puncta, reflecting a combination of antigen concentration and true synapse size, showed a small but significant reductions in the sizes of pan-Shank^+^ puncta on PTPσ-deficient neurons and GAD67-positive puncta on PTPδ-deficient neurons (**Fig 1A, 1B** and **Appendix Fig 2E**). Intriguingly, PTPσ KO, but not PTPδ KO, neurons exhibited marginally smaller cell body sizes (**Appendix Fig 2F, 2G, 2I** and **2J**), but similar dendritic branching as quantified by Sholl analysis, compared with control neurons (**Appendix Fig 2H** and **2K**). These results are consistent with the previously reported PTP KD effect (Han *et al*, 2018).

### Conditional PTPσ KO impairs excitatory synaptic transmission

To examine whether the reduced number of synapses in PTP-deficient neurons were accompanied by corresponding effects on the transmission of respective synapse types, hippocampal dissociated cultured neurons were assessed electrophysiologically (Fig 1D–1K). Lentivirus-mediated global loss of PTPσ specifically reduced the frequency and decay time, but not the amplitude, of excitatory synaptic transmission, as shown by measurement of miniature excitatory postsynaptic currents (mEPSCs) and miniature inhibitory postsynaptic currents (mIPSCs) (Fig 1D–1G). These results are consistent with the PTPσ KD effect on excitatory synaptic transmission (Dunah *et al*, 2005, Han *et al*, 2018, Ko *et al*, 2015). Moreover, lentivirus-mediated global loss of PTPδ had little effect on baseline synaptic transmission at both excitatory and inhibitory synapses (Fig 1H–1K). These results are different from the previous KD results showing specific reductions in excitatory (for PTPσ KD) and inhibitory (for PTPδ KD) synaptic transmission (Han *et al*, 2018), similar to the effects of gene manipulations on neuroligins (Chanda *et al*, 2017).

### Conditional PTPσ, but not PTPδ, KO alters synaptic vesicle tethering and active zone architectures

To further understand whether PTPσ-cKO affects synaptic structures, we performed transmission electron microscopy (TEM) with cultured cells. Presynaptic terminals and postsynaptic densities were examined and imaged from chemically fixed hippocampal neurons, as described (Acuna *et al*, 2016) (**Fig 2**). TEM analyses of cultured neurons showed that PTPσ-deficient and control presynaptic terminals contained similar numbers of total vesicles, but that the distribution of vesicles in the PTPσ-deficient terminals was significantly skewed (Fig 2A, 2D, 2E and 2F). Specifically, the number of membrane-proximal vesicles, defined as vesicles < 100 nm from the presynaptic AZ membrane, was significantly increased in PTPσ-deficient presynaptic terminals (**Fig 2E**). In contrast, the number of tethered vesicles, defined as vesicular structures docked at presynaptic AZ membranes, was only marginally increased, an increase that was not statistically significant (**Fig 2F**). Surprisingly, the AZ length was increased by ∼30% (**Fig 2A** and **2B**), similar to the doubled AZ size in *Caenorhabditis elegans* mutants lacking *ptp-3A*, an orthologue of the type IIa RPTP gene (Ackley *et al*, 2005, Han *et al*, 2019). Consistent with this, a corresponding increase (∼15%) in PSD length was observed (**Fig 2A** and **2C**). This phenotype was not observed in PTPδ-deficient presynaptic terminals, nor were there any alterations in other anatomical parameters (Fig 2H–2M). Quantitative immunoblot analysis of PTPσ- and PTPδ-deficient neurons showed comparable expression of presynaptic AZ and PSD proteins, although the level of RIM1/2 was significantly lower in PTPδ-deficient neurons (**Fig 2G, 2N, Appendix Fig 3A** and **3B**). These results suggest that PTPσ and PTPδ are not functionally redundant, and that PTPσ, but not PTPδ, is crucial in controlling the structural organization of both presynaptic AZs and PSDs.

**Figure 2.**
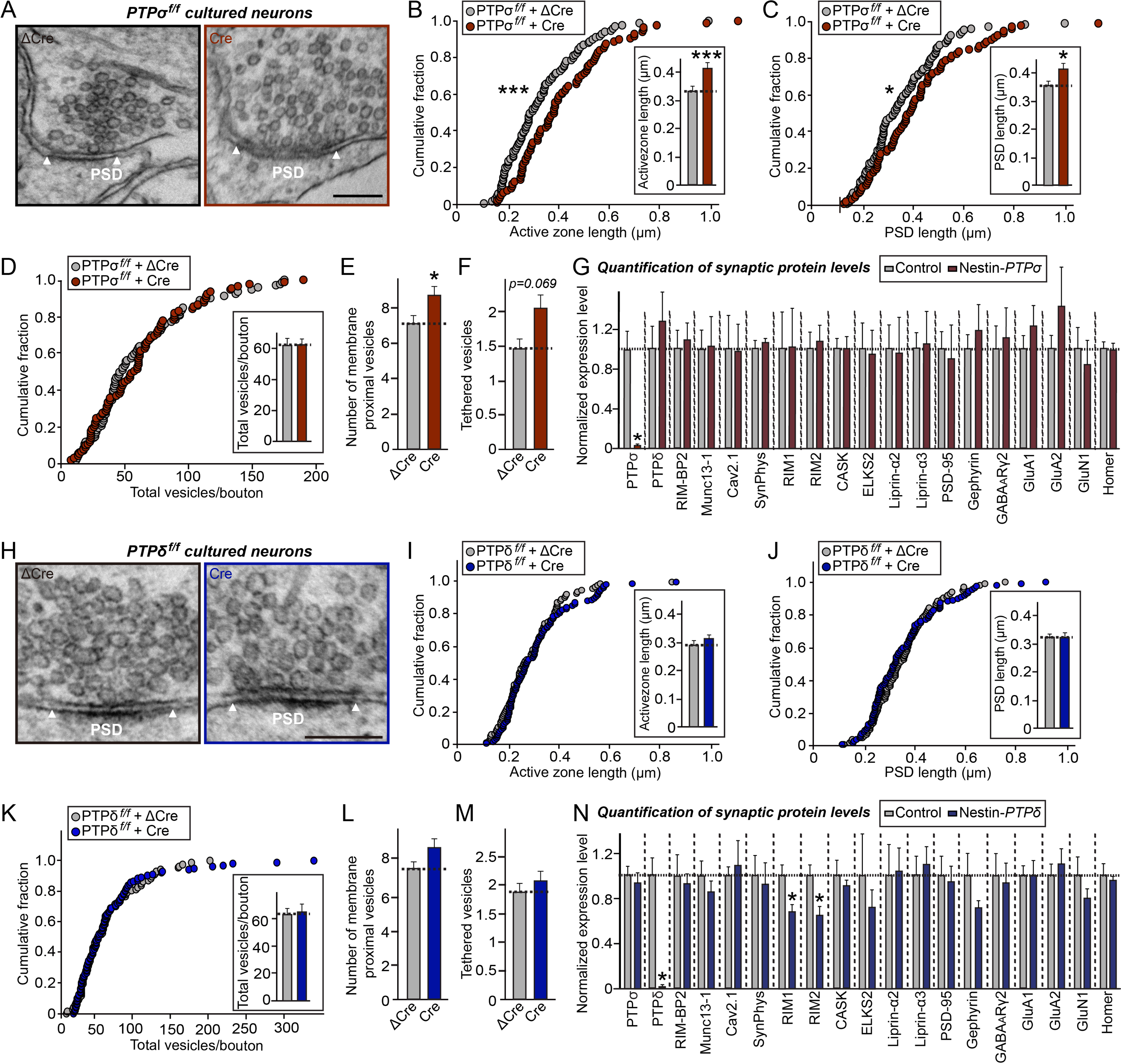
PTPσ deletion induces abnormal organization of synapse structures. **A** Representative electron micrographs of hippocampal neurons cultured from *PTPσ^f/f^* mice infected with lentiviruses expressing ΔCre (control) or Cre. **B, C** PTPσ deletion increases length of synaptic membranes. Cumulative distribution of the lengths of AZ (**B**) and PSD (**C**) for the indicated genotypes. Data are means ± SEMs (n denotes the number of analyzed neurons; ΔCre, 100 and Cre, 88; \**p* < 0.05; \*\*\**p* < 0.001; Mann Whitney U-test). **D–F** Total numbers of vesicles per bouton (**D**), membrane-proximal vesicles (**E**), and membrane-tethered vesicles (**F**) of control and PTPσ-deficient synapses. Data are means ± SEMs (n denotes the number of analyzed neurons; ΔCre, 100 and Cre, 88; \**p* < 0.05; Mann Whitney U-test). **G** Quantitative immunoblot analysis of PTPs, AZ proteins, and PSD proteins from control and Nestin-*PTPσ* mice. Data are means ± SEMs (n = 4 mice per group; \**p* < 0.05). **H** Representative electron micrographs of hippocampal neurons cultured from *PTPδ^f/f^* mice infected with lentiviruses expressing ΔCre (control) or Cre. **I, J** PTPδ deletion has no effect on the length of synaptic membranes. Cumulative distribution of the lengths of AZ (**I**) and PSD (**J**) for the indicated genotypes. Data are means ± SEMs (n denotes the number of analyzed neurons; ΔCre, 88 and Cre, 105). **K–M** Total number of vesicles per bouton (**K**), membrane-proximal vesicles (**L**), and membrane-tethered vesicles (**M**) of control and PTPδ-deficient synapses. Data are means ± SEMs (n denotes the number of analyzed neurons; ΔCre, 88 and Cre, 105). **N** Quantitative immunoblot analysis of PTPs, AZ proteins, and PSD proteins from control and Nestin-*PTPδ* mice. Levels of RIM1 and RIM2 expression were significantly lower in PTPδ-deficient than in control mice. Data are means ± SEMs (n = 4 mice per group; \**p* < 0.05; Mann Whitney U-test).

### Conditional KO of PTPσ reduces synaptic localization of excitatory synaptic vesicles in presynaptic boutons

Altered localization of synaptic vesicles in PTPσ-deficient nerve terminals may align with the possibility that PTPσ directly regulates the number of readily releasable vesicles (RRPs) (Zucker & Regehr, 2002). We thus examined whether the number of RRPs was changed at excitatory synapses by stimulating release of the entire RRP using a well-established hypertonic sucrose solution (500 mOsm) and quantifying RRP size by integrating the total charge transfer during the first 2 seconds of the release (Rosenmund & Stevens, 1996). Strikingly, the number of RRPs was significantly reduced in PTPσ-cKO neurons, as indicated by reductions of ∼14.7% in charge transfer and ∼31.2% in peak amplitude (**Fig 3A–3C**). This provides a hypothetical explanation for the positive regulation of neurotransmitter release by PTPσ at excitatory synapses, despite the increased anatomic proximity of synaptic vesicles to the active zone.

**Figure 3.**
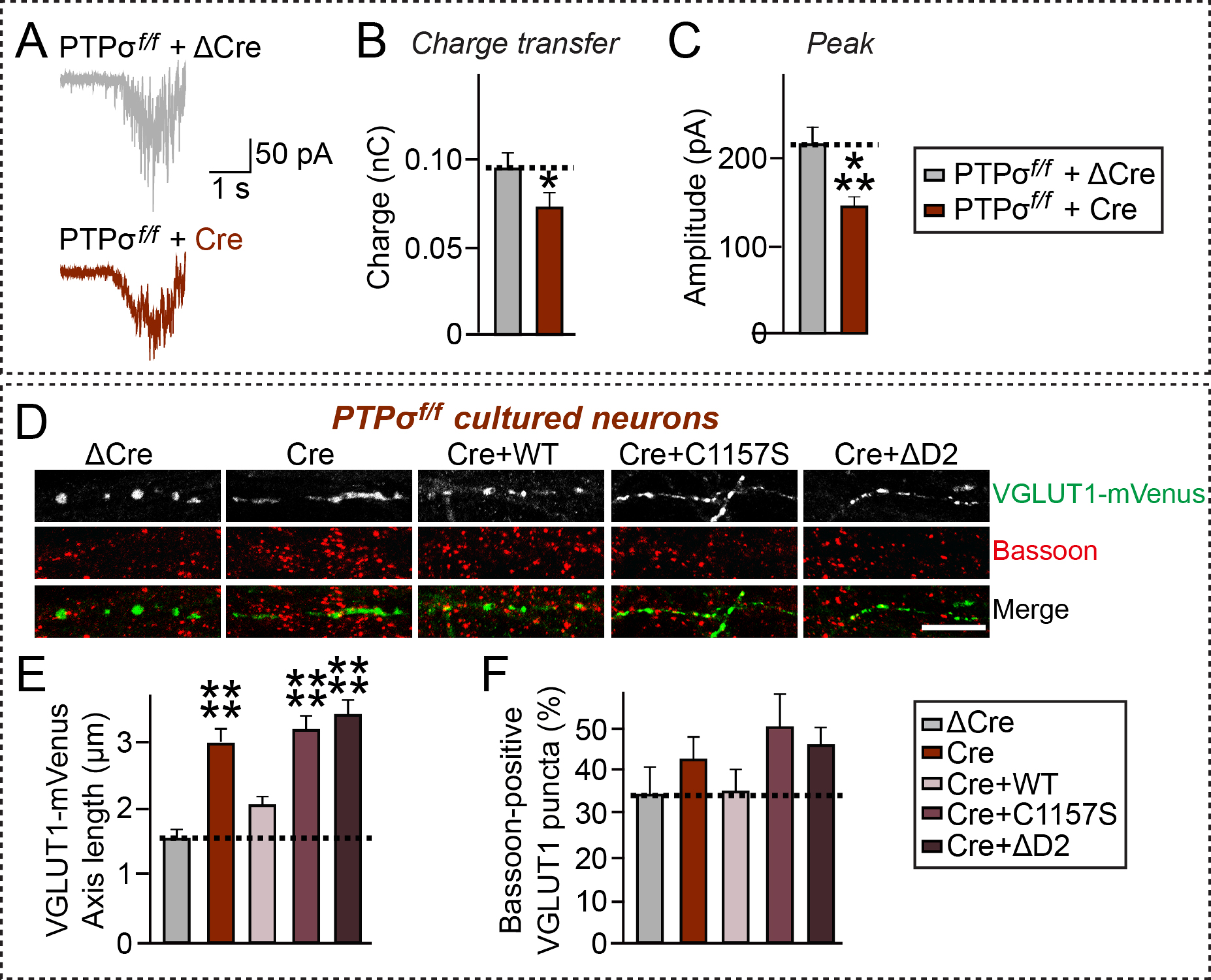
PTPσ deletion reduces vesicle localization in excitatory presynaptic boutons. **A** Representative traces of AMPAR-EPSCs evoked by single 2 sec pulse of 0.5 M sucrose delivered at a 1-min interval, recorded from hippocampal cultured neurons derived from *PTPσ^f/f^* mice infected with lentiviruses expressing inactive (ΔCre) or active (Cre) cre-recombinase. **B, C** Bar graphs showing charge transfer (**B**) and peak amplitudes (**C**) of sucrose-evoked EPSCs, estimated as the synaptic charge transfer integrated over 30 sec. Recordings were performed in the presence of 1 µM tetrodoxin and 50 µM picrotoxin. Data are means ± SEMs (n denotes number of analyzed neurons; ΔCre, 26 and Cre, 29; \**p* < 0.05; \*\*\**p* < 0.001). **D** Representative images of cultured neurons (DIV10) derived from *PTPσ^f/f^* mice infected with lentiviruses expressing ΔCre or Cre at DIV3 and transfected with VGLUT1-Venus (green) at DIV8. Anti-Bassoon (red) was used to mark the presynaptic active zone. Scale bar: 10 μm. **E** Quantification of synaptic vesicle diffusion from images in (**D**), determined by measuring the average length of the major axis of VGLUT1-Venus fluorescence in transfected axons. Data are means ± SEMs (n denotes the number of analyzed neurons; ΔCre, n = 19; Cre, n = 18; Cre+/PTPσ WT, n = 15; Cre+/PTPσ C1157S, n = 15; and Cre+/PTPσ ΔD2, n = 17; \*\*\*\**p* < 0.0001; ANOVA with a non-parametric Kruskal-Wallis test). **F** Quantification of VGLUT1-Venus fluorescence enrichment at presynaptic active zone for the images in (**D**). Data are means ± SEMs (n denotes the number of analyzed neurons; ΔCre, n = 19; Cre, n = 18; Cre+/PTPσ WT, n = 15; Cre+/PTPσ C1157S, n = 15; and Cre+/PTPσ ΔD2, n = 17; ANOVA with a non-parametric Kruskal-Wallis test).

To better understand the role of PTPs in organizing presynaptic functions at the excitatory synapse (see **Fig 1**), we infected cultured hippocampal PTPσ- or PTPδ-cKO neurons with ΔCre-EGFP or Cre-EGFP-expressing lentiviruses, transfected the neurons with venus-fused VGLUT1 (an excitatory synapse-specific vesicle marker) 5 days after the infections, and stained neurons with antibodies to Bassoon (a presynaptic active zone marker) 2 days after the transfections (**Fig 3**). We found that expression of VGLUT1-Venus in PTPσ-cKO neurons resulted in a more diffuse pattern of Venus-fused vesicular markers compared with control neurons infected with ΔCre-EGFP, indicating that ablation of PTPσ inhibits the synaptic localization of synaptic vesicles at excitatory presynaptic boutons (**Fig 3D** and **3E**). No changes in Bassoon localization at presynaptic boutons were observed in PTPσ-cKO neurons (**Fig 3D** and **3F**). To further dissect the mechanism by which PTPσ regulates vesicle localization at excitatory synapses, we designed three lentiviruses expressing PTPσ variants, based on validated HA epitope-tagged PTPσ variants that were previously used in cultured neurons (Han *et al*, 2018). These lentiviruses expressed PTPσ wild-type (WT), a PTPσ deletion mutant lacking the D2 domain (ΔD2), or a PTPσ point mutant defective in tyrosine phosphatase activity (C1157S). Lentiviral expression of PTPσ WT, but not other PTPσ variants, completely reversed the diffuse distribution pattern of VGLUT1-Venus fluorescence in PTPσ-cKO neurons, producing a punctate pattern (**Fig 3D** and **3E**). In addition, synaptic localization of vesicles was not rescued by expression of PTPσ intracellular mutants, suggesting that PTPσ requires D2 domain-mediated molecular interactions and tyrosine phosphatase activity to appropriately direct excitatory synaptic vesicles into presynaptic boutons. Collectively, these results suggest that PTPσ is specifically involved in presynaptic assembly by organizing vesicle localization at excitatory synapses using intracellular mechanisms.

### Conditional PTPσ KO in mPFC and CA1 neurons reduces excitatory presynaptic innervation onto postsynaptic target neurons and excitatory neurotransmitter release

Although the synaptic roles of invertebrate orthologues of type IIa RPTPs have been primarily studied from the perspective of their presynaptic structure and function (Chagnon *et al*, 2004, Um & Ko, 2013), mammalian PTPσ and PTPδ appear to be expressed in both presynaptic and postsynaptic neurons (Dunah *et al*, 2005, Han *et al*, 2018) (**Appendix Fig 3C**). Because the functional locus of a specific synaptic protein cannot be precisely determined in cultured neurons, a Cre driver line under the control of a Wolfram syndrome 1 homolog (Wfs1) promoter was utilized (Kitamura *et al*, 2014, Madisen *et al*, 2010). The presence of Wfs1-positive neurons, including in the dorsal CA1 and layer II/III of the mPFC, were confirmed by robust tdTomato expression in the Ai9 reporter mouse line (Luuk *et al*, 2008, Madisen *et al*, 2010) (**Appendix Fig 4A**). Immunohistochemical analysis of Wfs1 expression in the mPFC and hippocampal CA1 showed strong Wfs1-immunoreactive signals in Tbr1-positive excitatory neurons but not GAD67-positive GABAergic neurons (**Appendix Fig 4B**). Because presynaptic organization was abnormally altered in PTPσ-deficient cultured neurons (**Fig 2**), the pre- and postsynaptic effects of PTPσ deletions were analyzed by focusing on presynaptic CA1 neurons of the hippocampus at synapses formed onto postsynaptic pyramidal neurons in the subiculum and on those in presynaptic layer II/III neurons at synapses formed onto postsynaptic layer V neurons of the mPFC (**Figs 4** and **5**). *PTPσ*^f/f^ mice were crossed with a Wfs1-Cre driver line to yield Wfs1-*PTPσ* mice, which were viable and fertile and comparable in size to control mice (**Appendix Fig 4C**). Moreover, NeuN and Nissl staining of Wfs1-*PTPσ* brains showed normal gross morphology (**Appendix Fig 4D** and **4E**). Anatomical changes at synapses formed by presynaptic CA1 region neurons on postsynaptic subicular neurons and presynaptic layer II/III layer neurons on postsynaptic layer V neurons were evaluated by quantitative immunofluorescence analyses (**Fig 4A**), which showed that the density and integrated intensity of VGLUT1 puncta were significantly reduced in subicular, but not mPFC layer V, neurons (**Fig 4B–4F**). However, the density and intensity of GAD67 puncta in the corresponding brain regions were comparable in Wfs1-*PTPσ* and control mice (**Fig 4G–4K**). Adeno-associated viruses (AAVs) expressing Cre-recombinase (AAV-Cre) or inactive Cre-recombinase (AAV-ΔCre) were stereotactically injected into ventral hippocampal CA1 (vCA1) of *PTPσ*^f/f^ and *PTPδ*^f/f^ mice. Subsequent quantitative immunohistochemical analyses showed decreased excitatory (but not GABAergic) innervations onto subicular neurons from *PTPσ*^f/f^ mice infected with AAV-Cre, and decreased GABAergic (but not excitatory) innervation onto subicular neurons from *PTPδ*^f/f^ mice infected with AAV-Cre (**Fig EV2**).

**Figure 4.**
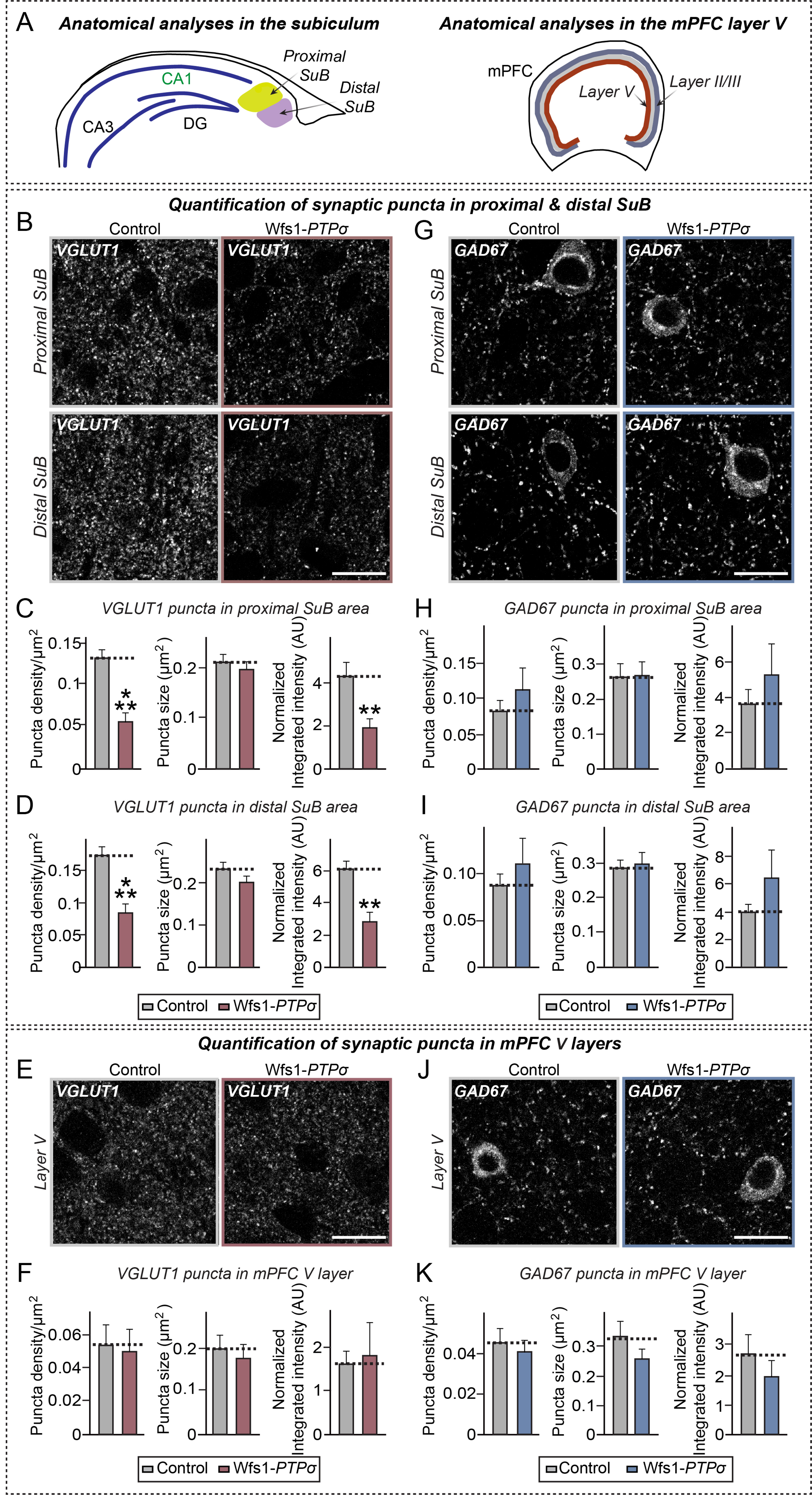
Wfs1-*PTPσ* KO mice exhibit decreased excitatory synaptic innervation in postsynaptic target neurons. **A** Schematic depiction of anatomical analyses in the subiculum (SuB; left) and the medial prefrontal cortex (mPFC; right). Each subiculum was divided into the proximal and distal subiculum. **B, G** Representative immunofluorescence images of the proximal and distal SuB of Ctrl and Wfs1-*PTPσ* mice using VGLUT1 (**B**) or GAD67 (**G**). Scale bar: 20 μm. **C, H** Quantification of the density, size and integrated intensity of VGLUT1-positive (**C**) and GAD67-positive (**H**) synaptic puncta in the proximal SuB. Data are means ± SEMs (n denotes the number of analyzed brain mice; 8 mice per each group; \*\**p* < 0.01 and \*\*\**p* < 0.001; Mann Whitney U-test). **D, I** Quantification of the density, size and integrated intensity of VGLUT1-positive (**D**) and GAD67-positive (**I**) synaptic puncta in the distal SuB. Data are means ± SEMs (n denotes the number of analyzed mice; 8 mice per each group; \*\**p* < 0.01, \*\*\**p* < 0.001; Mann Whitney U-test). **E, J** Representative immunofluorescence images of the mPFC layer V of Ctrl and Wfs1-*PTPσ* mice using VGLUT1 (**E**) or GAD67 (**J**). Scale bar: 20 μm. **F, K** Quantification of the density, size and integrated intensity of VGLUT1-positive (**F**) and GAD67-positive (**K**) synaptic puncta in the mPFC layer V. Data are means ± SEMs (n denotes the number of analyzed brain mice; 8 mice per each group; Mann Whitney U-test).

**Figure 5.**
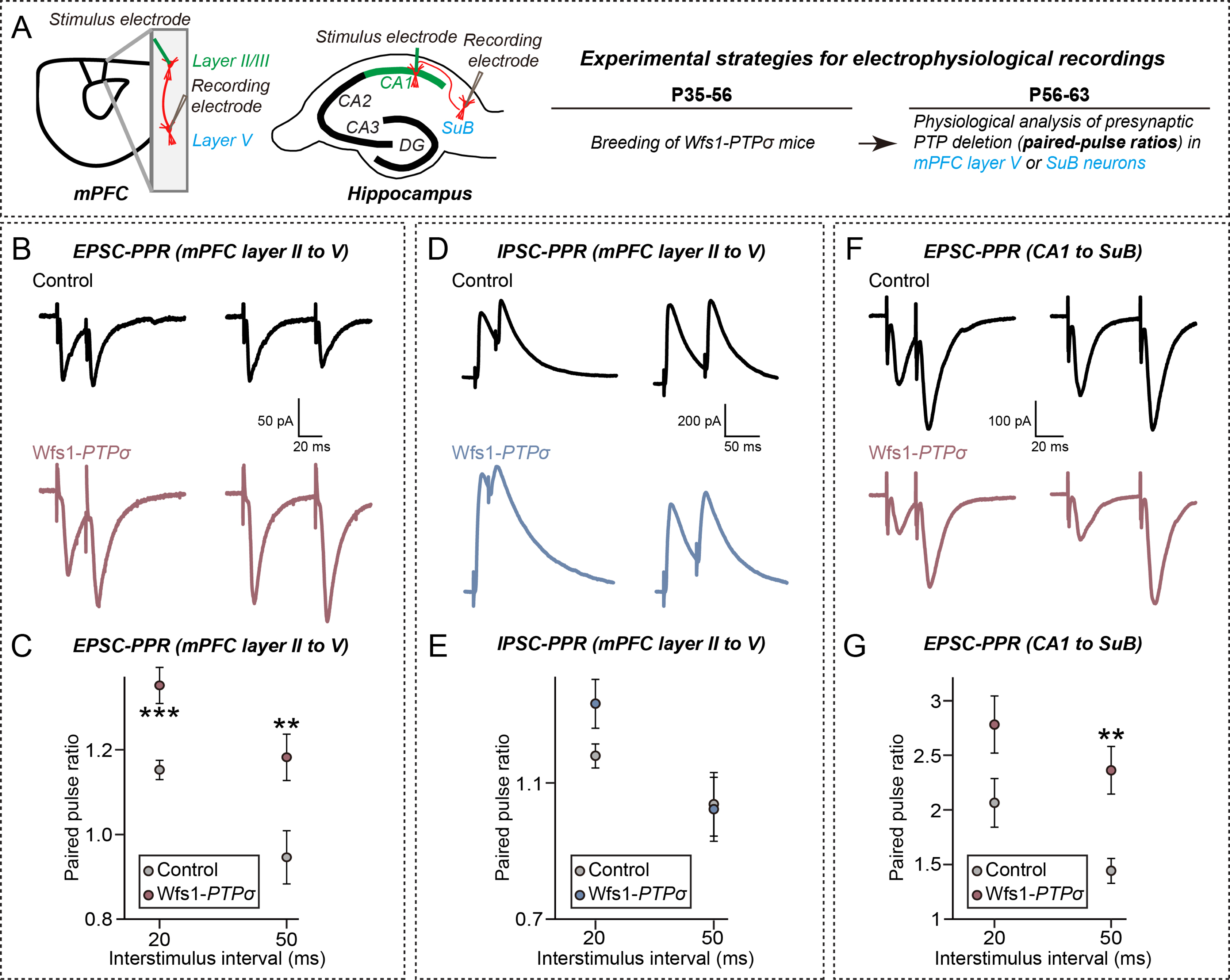
Presynaptic deletion of PTPσ, but not of PTPδ, impairs neurotransmitter release at excitatory synapses of postsynaptic target neurons. **A** Experimental strategies for electrophysiological recordings in mPFC and hippocampal SuB neurons of Wfs1-*PTPσ* mice. **B, D** Representative traces of paired pulse ratios (PPRs) of EPSCs (**B**) and IPSCs (**D**) in synapses of mPFC layer II to layer V at two different interstimulus intervals (20 and 50 ms for EPSC-PPRs and 20 and 50 ms for IPSC-PPRs). **C, E** EPSC-PPRs (**C**) and IPSC-PPRs (**E**) in synapses of mPFC layers II–V as a function of the indicated interstimulus intervals. Data are means ± SEMs (\*\**p* < 0.01, \*\*\**p* < 0.001; two-tailed Student’s t-test). **F** Representative traces of paired pulse ratios (PPRs) of EPSCs in hippocampal CA1-SuB synapses at two different interstimulus intervals (20 and 50 ms). **G** EPSC-PPRs in hippocampal CA1-SuB synapses as a function of the indicated interstimulus intervals. Data are means ± SEMs (\*\**p* < 0.01; two-tailed student’s t-test).

To further corroborate these anatomical observations and identify any possible presynaptic changes, PPRs in subicular and mPFC layer V neurons were measured (**Fig 5A**). Although EPSC-PPRs were significantly increased in both synapses (Fig 5B, 5C, 5F and 5G), IPSC-PPRs were not changed in mPFC layer V neurons from Wfs1-*PTPσ* mice (Fig 5D and 5E). Taken together, these results suggest that PTPσ is a critical modulator of presynaptic innervations and neurotransmitter release at excitatory synapses.

### Conditional PTPσ deletions exert no postsynaptic effect

Next, the effect of presynaptic deletion of PTPσ on basal synaptic transmission was analyzed in Wfs1-*PTPσ* mice. Surprisingly, there was a marginal reduction in frequency, but not amplitude, of spontaneous EPSCs (sEPSCs) in mPFC layer V (but not subicular pyramidal) neurons (**Fig EV3A–EV3C** and **EV3G–EV3I**), but no detectable effect on the frequency or amplitude of mIPSCs in mPFC layer V pyramidal neurons (**Fig EV3D–EV3F**). The ratio of NMDA-to AMPA-receptor mediated EPSCs (i.e. NMDA/AMPA ratio) was assessed by stimulating Schaffer collateral axons of hippocampal CA3 neurons or axons of hippocampal CA1 neurons, and measuring postsynaptic responses in hippocampal CA1 neurons and subicular neurons, respectively (**Fig 6D, 6E** and **Appendix Fig 5**). In agreement with findings showing no prominent phenotype affecting basal excitatory synaptic transmission in neurons lacking PTPσ, there were no changes in the density of excitatory and inhibitory synapses, in NMDA/AMPA ratio, and in the frequency and amplitude of excitatory synaptic transmission in PTPσ-deficient CA1 pyramidal neurons (**Fig 6** and **EV3J–EV3L**). These results suggest that PTPσ primarily functions presynaptically and does not *trans*-synaptically regulate postsynaptic responses *in vivo*.

**Figure 6.**
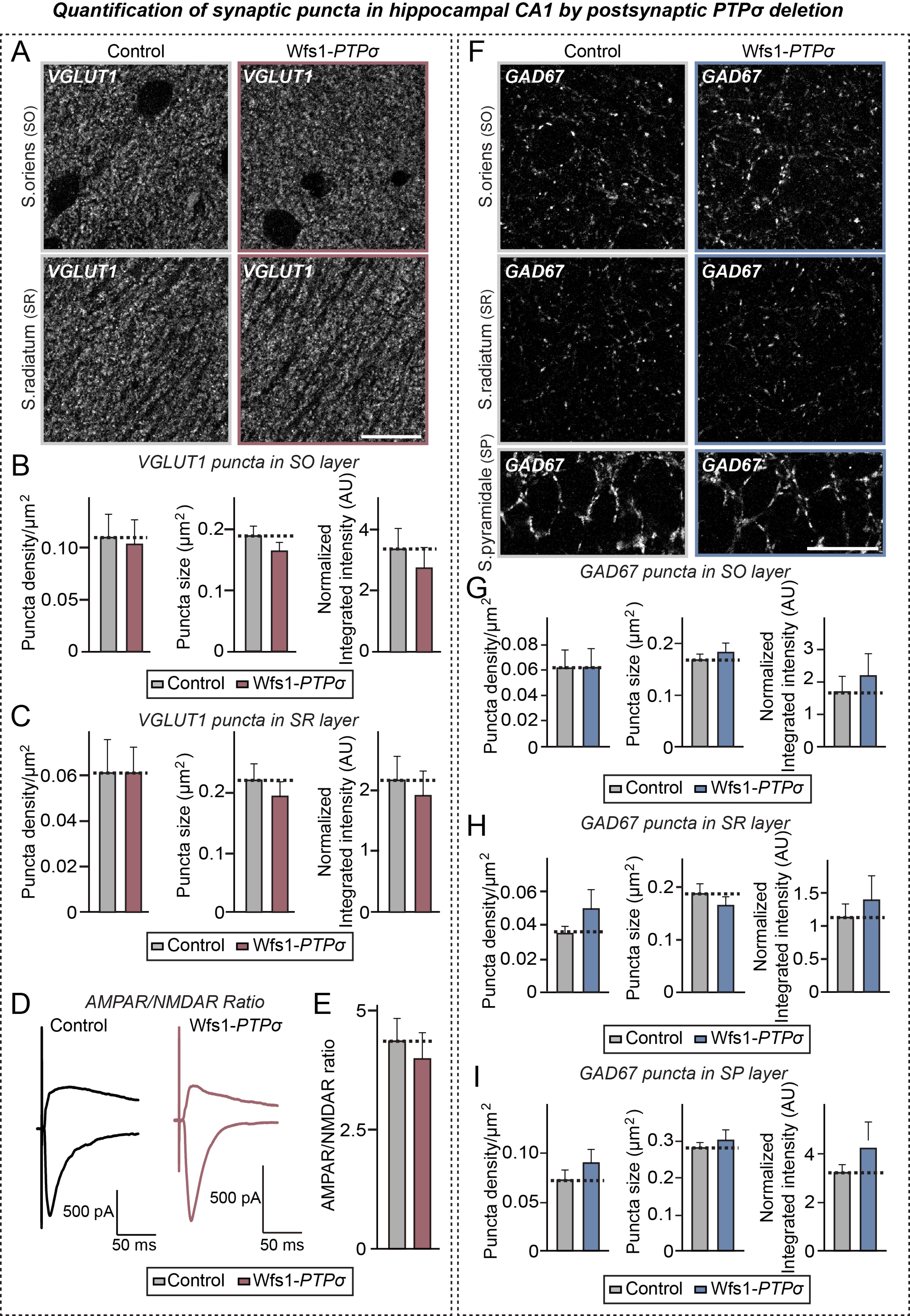
Postsynaptic deletion of PTPσ exerts no effects on excitatory synapse organization in hippocampal CA1 region. **A, F** Representative immunofluorescence images of the *stratum radiatum* (SR) and *stratum oriens* (SO) layer of control and Wfs1-*PTPσ* mice using VGLUT1 (**A**) or GAD67 (**F**). Scale bar: 20 μm. **B, C** Quantification of the density, size and integrated intensity of VGLUT1-positive synaptic puncta in the SO (**B**) and SR (**C**) layers of the hippocampal CA1 region. Data are means ± SEMs (n = 8 mice per each group; Mann Whitney U-test). **D, E** Representative traces (**D**) of evoked EPSCs at holding potentials of −70 mV and +40 mV in control and Wfs1-*PTPσ* mice. (**E**) AMPAR/NMDAR ratios, calculated by dividing the EPSC peak amplitude at 10 ms (−70 mV) by the EPSC amplitude at 130 ms (+ 40 mV). Data are means ± SEMs (n = 14 cells from 4–5 mice; two-tailed student’s t-test). **G–I** Quantification of the density, size and integrated intensity of GAD67-positive synaptic puncta in the SO (**G**), SR (**H**) and SP (**I**) layers of the hippocampal CA1 region. Data are means ± SEM (n = 8 mice per each group; Mann Whitney U-test).

### PTPσ controls formation of excitatory inputs via intracellular mechanisms

To interrogate whether interactions of PTPσ with intracellular scaffolds and/or substrates are responsible for presynaptic innervation at excitatory synapses, we designed AAVs expressing PTPσ WT and its intracellular variants (ΔD2 and C1157S) (**Fig EV4A**). PTPσ expression was driven by the neuron-specific synapsin promoter, and immunohistochemically with anti-HA antibody (**Fig EV4**). These PTPσ AAVs, along with AAV-Cre, were stereotactically injected into the vCA1 of adult male Wfs1-*PTPσ* mice, with quantitative immunofluorescence analyses performed after 3 weeks after stereotactic surgery. Expression of WT PTPσ, but not the other PTPσ variants, significantly increased the density of VGLUT1 puncta in proximal subiculum regions of Wfs1-*PTPσ* mice (**Fig EV4**). The density of VGLUT1 puncta in distal subiculum regions was not influenced, in line with the specific connectivity between the vCA1 and proximal subiculum (O’Mara, 2005). Expression of PTPσ intracellular mutants failed to completely reverse the deficits in innervation of excitatory synaptic inputs (**Fig EV4**). These results suggest that PTPσ requires intracellular mechanisms (D2-mediated interactions and tyrosine phosphorylation) to organize innervation of excitatory inputs onto postsynaptic targets *in vivo*.

### PTPσ deletion in ventral CA1 induces abnormal anxiety-like behavior

To assess the behavioral consequences of the PTPσ deletion in mice, adult (9–11-week old) male Wfs1-*PTPσ* mice and their control littermates were screened using a behavioral panel designed to assess locomotion, anxiety, sociability, and learning (**Fig 7** and **EV5**). In the Laboras test, in which mouse movements were recorded continuously for 48 hours, both control and Wfs1-*PTPσ* mice exhibited overall similar home cage-like activities, although Wfs1-*PTPσ* mice showed a marginal reduction in the amount of food eaten (**Fig EV5A** and **EV5B**). Wfs1-*PTPσ* mice did not show anxiety-like behavior in open-field tests, spending an amount of time in the center region of the open-field arena (**Fig EV5C** and **EV5D**), and in light-dark transition tests, as measured by time spent in the light box and number of total entries in the light-dark apparatus (**Fig EV5E** and **EV5F**). However, these mice were less anxious in the elevated plus-maze (EPM) test, spending more time in open arms (**Fig 7A–7C**), suggesting that deletion of PTPσ from discrete brain regions manifests as anxiolytic-like behaviors under specific conditions. In tests for various forms of memory, Wfs1-*PTPσ* mice exhibited no abnormalities in Y-maze tests, which assess working memory (**Fig EV5G** and **EV5H**), and novel object-recognition tests, which assess recognition memory (**Fig EV5I** and **EV5J**). Meanwhile, Wfs1-*PTPσ* mice exhibited low freezing time in contextual fear conditioning tests, which assess fear memory (**Fig EV5K** and **EV5L**). Intriguingly, analysis of two different facets of social behaviors in three-chamber tests revealed that Wfs1-*PTPσ* mice exhibit mildly impaired sociability, spending less time in the chamber containing the stranger mouse than in either of the empty chambers (**Fig 7D–7F**). However, when the inanimate object was replaced by another stranger mouse, both control and Wfs1-*PTPσ* mice showed comparable preference for exploring the new stranger mouse (**Fig 7D–7F**). These results suggest that Wfs1-*PTPσ* mice are impaired in social interactions (sociability), whereas social novelty recognition and social anxiety remained intact.

**Figure 7.**
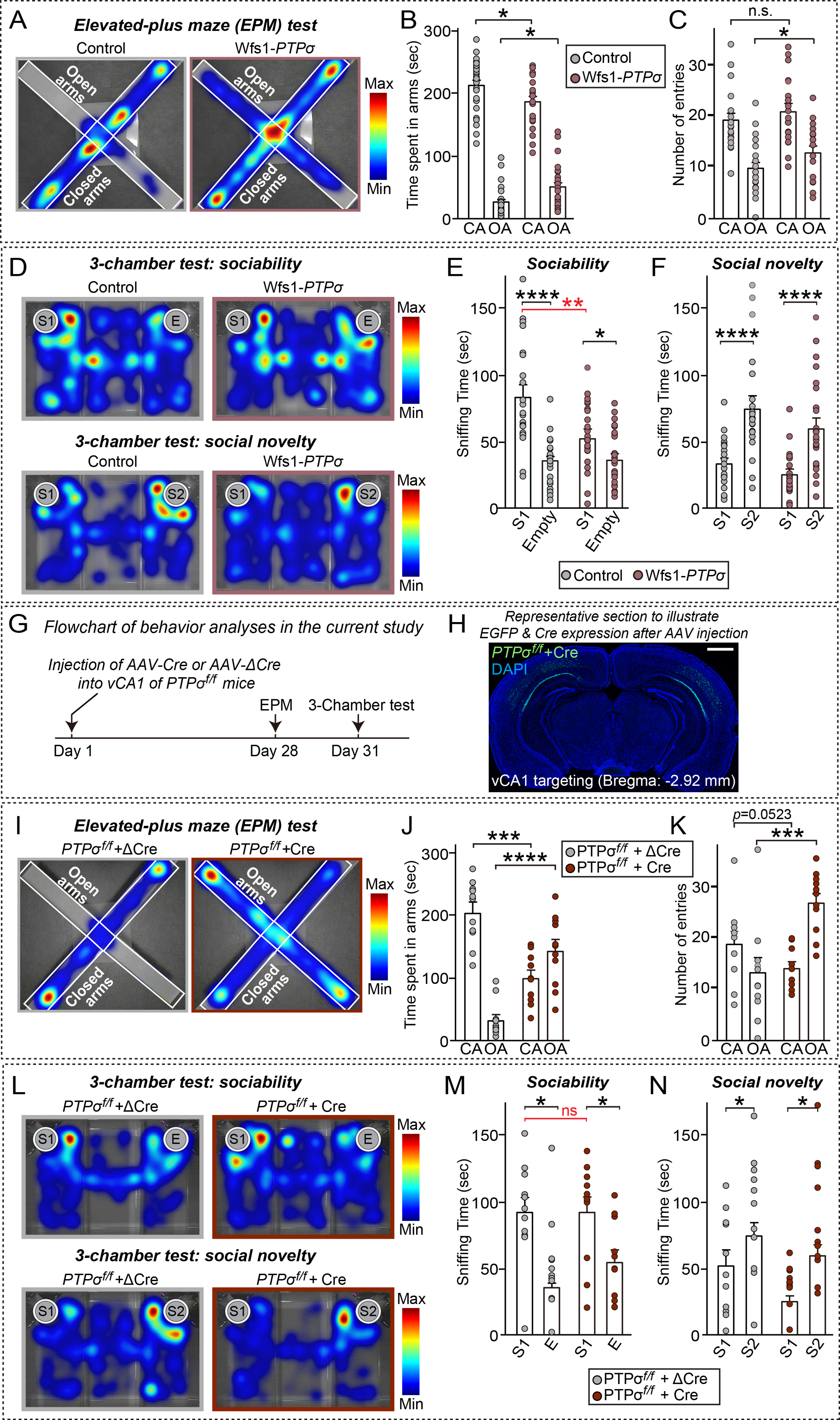
Presynaptic deletion of PTPσ in ventral hippocampal CA1 selectively induces abnormal anxiety-like behaviour. **A–C** Analysis of control and Wfs1-*PTPσ* mice in the elevated-plus maze (EPM) tests. Representative heat map (**A**) and graphs showing time spent in open arms (**B**) and number of entries (**C**). Data are means ± SEMs (control, n=24; Wfs1-*PTPσ,* n=23; \**p* < 0.05; Mann Whitney U-test). **D–F** Analysis of control and Wfs1-*PTPσ* mice in three-chamber tests. Representative heat map (**D**) and quantification of sociability (**E**) and social novelty (**F**). Sniffing time was defined as the time spent sniffing the novel mouse and the novel object. Abbreviations: E, empty cup; S1: stranger mouse 1; S2: stranger mouse 2. Data are means ± SEMs (control, n = 21; and Wfs1-*PTPσ,* n = 24; \**p*< 0.05, \*\**p*< 0.01, \*\*\*\**p* < 0.0001; Mann Whitney U-test). **G** Schematic diagram of mouse behavior. AAV-ΔCre or AAV-Cre was bilaterally injected into the ventral CA1 (vCA1) region of the hippocampus of ∼6 week-old PTPσ^f/f^ mice. Four weeks later, these mice were subjected to the EPM and three-chamber tests in the indicated order. Abbreviation: EPM, elevated plus-maze test. **H** Representative coronal section showing EGFP expression after AAV injection the vCA1. Scale bar, 1 mm. **I–K** Analysis of ΔCre- and Cre-injected PTPσ^f/f^ mice in the elevated-plus maze (EPM) tests. Representative heat map (**I**) and graphs showing time spent in open arms (**J**) and number of entries (**K**). Data are means ± SEMs (PTPσ *^f/f^* + ΔCre, n = 11; and PTPσ *^f/f^* + Cre, n = 11; \*\*\**p* < 0.001, \*\*\*\**p* < 0.0001; Mann Whitney U-test). **L–N** Analysis of ΔCre- and Cre-injected PTPσ^f/f^ mice in three-chamber tests. Representative heat map (L) and quantification of sociability (**M**) and social novelty (**N**). Sniffing time was defined as time spent sniffing the novel mouse and the novel object. Abbreviations: E, empty cup; S1: stranger mouse 1; S2: stranger mouse 2. Data are means ± SEMs (PTPσ *^f/f^* + ΔCre, n = 11; and PTPσ *^f/f^* + Cre, n = 11 mice; \**p* < 0.05; Mann Whitney U-test).

To identify the specific brain regions responsible for two prominent behavioral phenotypes, anxiety-like behavior and sociability, one of the brain regions (vCA1), in which Wfs1 promoter activity was positive, was selected (**Appendix Fig 4A**). The vCA1 neurons of adult male *PTPσ*^f/f^ mice were stereotactically injected with AAV-EGFP/ΔCre (ΔCre) or AAV-EGFP/Cre (Cre) separately (**Fig 7G** and **7H**), and the mice were subjected to the EPM and three-chamber tests (**Fig 7I–7N**). Surprisingly, we found that Cre expression in the vCA1 region replicated the impairment in anxiety-like behavior of Wfs1-*PTPσ* mice in the EPM test, but had no effect on the three-chamber test (**Fig 7I–7N**). These results suggest that PTPσ presynaptically regulates the operation of a specific neural circuit that mediates distinct mouse behavior.

## Discussion

Although many studies have unequivocally shown that Nrxns are crucial at synapses, few studies have evaluated the *in vivo* roles of vertebrate LAR-RPTPs. Investigations using constitutive KO mice lacking one or two LAR-RPTPs, or shRNA-mediated KD approaches, showed that LAR-RPTPs are significant regulators of various aspects of nervous system development (Chagnon *et al*, 2004, Han *et al*, 2016, Han *et al*, 2019, Takahashi & Craig, 2013, Um & Ko, 2013). These studies showed that LAR-RPTPs affect the development and/or maturation of synapses as well as having potential compensatory adaptations and possibly unintentional off-target phenomena during development. Therefore, the present study involved the generation of PTPσ and PTPδ floxed mice, with these conditional KO mice analyzed to determine the synaptic roles of PTPσ and PTPδ. To this end, we employed a variety of experimental approaches including high-density cultured neurons generated using Cre-expressing lentiviruses *in vitro*, acute slices and tissues obtained from mice crossed with Cre driver lines, and slices and tissues obtained from3 mice stereotactically injected *in vivo* with Cre-expressing AAVs into specific brain areas (i.e. vCA1).

Our *in vitro* data demonstrated that PTPσ KO, but not PTPδ KO, specifically reduced the numbers of excitatory synapses and basal excitatory synaptic transmission, in agreement with previous KD studies (Han *et al*, 2018, Ko *et al*, 2015) (**Fig 1**). Moreover, PTPσ KO altered vesicle tethering and ultrastructural features (**Fig 2**), decreased RRP size and synaptic localization of excitatory synaptic vesicles (**Fig 3**). Our *in vivo* results further suggest that PTPσ presynaptically functions at two different types of synapses involving layer II–V connections in mPFC and CA1–subiculum connection in the hippocampus by modulating glutamate release (**Figs 4–6**), and that this regulation guides a specific behavior (**Fig 7**). Compared with the pervasive loss-of-function consequences of Nrxns in various cell-types across diverse brain areas and neural circuits (Südhof, 2017), the effects of conditional deletion of PTPσ were relatively marginal, whereas the effects of conditional deletion of PTPδ were surprisingly subtle or unnoticeable. These findings were unexpected, as members of the LAR-RPTP family bind to various synapse organizers crucial for discrete aspects of synapse development (Südhof, 2017, Um & Ko, 2013). Nevertheless, the current study revealed that vertebrate LAR-RPTPs perform unexpected distinct functions, which may help shape specific synaptic properties across many overlapping neural circuits.

### Dissection of PTPσ *in vivo* functions using sophisticated approaches

A rich body of previous studies have shown that invertebrate LAR-RPTPs have presynaptic roles in controlling AZ assembly and proper vesicle localization (Han *et al*, 2019). A few studies of vertebrate LAR-RPTP function have revealed their critical roles in postsynaptic neurons, including their regulation of spine morphogenesis development and their stabilization of surface AMPA-type glutamate receptors (Dunah *et al*, 2005, Wyszynski *et al*, 2002). Because several LAR-RPTP ligands have been identified as putative postsynaptic organizers in rodent neurons, LAR-RPTPs were thought to act in presynaptic neurons in a manner similar to Nrxns (Südhof, 2012). Consistent with previous findings (reviewed in Han *et al*, 2019), the results in the current study showed the loss-of-function phenotype of PTPσ, demonstrating that conditional KO of PTPσ in cultured hippocampal neurons specifically impairs excitatory, but not inhibitory, synaptic transmission (**Fig 1**). In contrast, conditional KO of PTPδ did not significantly affect most synaptic parameters (**Figs 1** and 2). These results were quite surprising and unexpected, as PTPδ is strongly expressed in brains and constitutive PTPδ KO was found to impair learning, alter long-term synaptic plasticity, and cause abnormal dendritic arborization (Nakamura *et al*, 2017, Uetani *et al*, 2000). Moreover, PTPδ KD produced pleiotropic effects on both presynaptic and postsynaptic processes (Dunah *et al*, 2005, Han *et al*, 2018, Li *et al*, 2015, Yim *et al*, 2013). Some discrepancies between the results of our and previous studies may be due to differences in developmental changes caused by constitutive KO and the well-documented off-target effects of RNA interference. Secondary effects triggered by KD may also have contributed to previously observed severe phenotypes; for example, PTPδ KD induced PTPσ upregulation in neurons, and vice versa, whereas PTPδ KO did not (Han *et al*, 2018) (**Fig 2N** and **Appendix Fig 3B**). Nevertheless, PTPσ KO and PTPσ KD had similar overall effects on synapse density and transmission *in vitro*, indicating the importance of confirmatory analyses using a sophisticated system and approaches in elucidating PTPσ function *in vivo*. It is possible, however, that the approaches employed in this study were not sufficiently sensitive to detect subtle changes in synapse properties. Moreover, the selection of experimental preparations may preclude detection of synaptic roles of PTPδ at *in vivo* synapses. In addition, PTPσ may functionally compensate for PTPδ loss, but not vice versa, whereas PTPδ may have peripheral roles in the operation of specific neural circuits in which, PTPσ is expressed. The use of cKO models to study the canonical and non-canonical roles of PTPσ and PTPδ *in vivo* at various stages of synapse development in other classes of neurons is warranted and may provide a more sophisticated understanding of how LAR-RPTPs act as a multivalent signaling platform in presynaptic neurons.

### Function of PTPσ as a presynaptic adhesion molecule in regulating excitatory synapse properties at mammalian excitatory synapses

We found that loss of PTPσ resulted in increased length of the presynaptic AZ and PSD membranes, and increased numbers of membrane-proximal vesicles (**Fig 2**). Some of these phenotypes were similar to those of KO mice lacking a subset of AZ proteins involved in vesicle tethering and recruitment of VGCCs to AZ membranes, and were also similar to loss-of-function phenotypes of *Syd-2* mutations in *C. elegans* (an orthologue of liprin-α) (Acuna *et al*, 2016, Dai *et al*, 2006, Grauel *et al*, 2016, Kaeser *et al*, 2011, Wang *et al*, 2016, Zhen & Jin, 1999). Although a subset of these phenotypes may reflect adaptive changes as a homeostatic mechanism, PTPσ likely contributes to the normal distribution of SVs and the organization of *trans*-synaptic nanomolecular complexes for efficient glutamate release (**Fig 2**). Importantly, investigation of common/redundant functions of LAR-RPTPs using conditional triple KOs of PTPσ, PTPδ, and LAR is required to provide further insights into the major observations of the current study.

Impaired glutamate release efficiency observed in PTPσ cKO mice has been observed in several prior studies reporting loss-of-function of Nrxn1α (Etherton *et al*, 2009), liprin-α2 (Spangler *et al*, 2013), and RIM1 (Kaeser *et al*, 2008). Strikingly, PTPσ KO induces abnormal vesicle positioning close to AZ membranes (**Fig 2**), but drastically suppressed RRP size in cultured neurons (**Fig 3**) that is controlled by positional priming (Neher & Sakaba, 2008). These results suggest that PTPσ KO may disorganize AZ structure and function by decoupling key nanomolecular machinery in AZs.

Our genetic manipulations using the Wfs1-Cre driver line allowed determination of the presynaptic and postsynaptic role of PTPσ in excitatory neurons (**Figs 4**–**7**). PTPσ deletion in mPFC layer II or hippocampal CA1 reduced structural innervation of excitatory inputs and increased EPSC-PPRs in mPFC layer V or subicular neurons, along with a marginal reduction in frequency, but not amplitude, of sEPSCs (**Fig 4, 5** and **EV3**). Intriguingly, PTPσ deletion in hippocampal CA1 failed to elicit any changes in Schaffer collateral evoked excitatory synaptic transmission or basal synaptic transmission (**Fig 6**), indicating that postsynaptic strength is not *trans*-synaptically regulated by PTPσ *in vivo*. These results suggest that PTPσ primarily functions as a presynaptic adhesion molecule to regulate the amount and/or pattern of excitatory inputs, resulting in the modulation of neurotransmitter release.

### PTPσ requires various molecular mechanisms to regulate excitatory neurotransmitter release

PTPσ employs various extracellular and intracellular mechanisms in mediating excitatory presynaptic assembly (Han *et al*, 2018). Utilization of *in vivo* replacement strategies in PTPσ cKO mice revealed that multiple molecular mechanisms contribute to the regulation of innervation of excitatory synaptic inputs of CA1 neurons onto subiculum neurons (**Fig EV4**). These findings suggest that the ability of PTPσ to regulate different presynaptic properties is due to the concerted actions of various mechanisms—including the availability of release-ready SVs, the integrity of AZ organization, and the magnitude of Ca^2+^ influx—by coordinating extracellular signals with intracellular machineries responsible for AP-gated glutamate release of SVs. The most parsimonious mechanism by which PTPσ likely regulates glutamate release is through the direct interaction of PTPσ with liprin-α via its intracellular D2 domain. Liprin-α3 functions as an upstream AZ scaffolding factor in AZ assembly, further interacting with RIMs and other presynaptic proteins, including ELKS, mDiaphanous, CASK, and GIT1 (Han *et al*, 2019, Südhof, 2012, Wong *et al*, 2018). Because PTPσ requires intracellular complexes for presynaptic assembly (Han *et a*l, 2018), PTPσ may mediate various presynaptic functions by coordinating the interactions of liprin-α with other presynaptic proteins. Alternatively, a subset of PTPσ substrates may be involved in PTPσ-mediated glutamate release. Several validated PTPσ substrates have been associated with the regulation of actin cytoskeleton remodeling processes (Han *et al*, 2018, Um & Ko, 2013). Systematic analyses using cKO mice lacking PTPσ substrates, combined with a high-resolution approach to identify putative substrates, are crucial to fully understand how a network of actin filaments associated with SVs is coordinated with neurotransmitter release (Doussau & Augustine, 2000, Nelson *et al*, 2013). The ability of PTPσ, along with specific Cav2 complexes, to directly regulate Ca^2+^ currents should also be rigorously examined. Extracellular interactions of PTPσ with postsynaptic ligands could control the function of VGCCs (Muller *et al*, 2010). This hypothesis was supported by a study showing that KD of Slitrk1, a postsynaptic ligand for PTPσ, increased the number of docked SVs, mimicking some of the loss-of-function phenotype of PTPσ cKO (Schroeder *et al*, 2018) (**Fig 3**).

### Role of PTPσ function in neural circuits: canonical or context-specific function

One prominent mechanism of altered neurotransmitter release in regulating specific properties of neural circuits is STP. STP has been implicated in many cognitive functions involving mPFC and the hippocampus, including short-term working memory, information processing, and decision-making processes (Abbott & Regehr, 2004, Mongillo *et al*, 2008). Because this form of plasticity was thought to reflect a presynaptic change in Ca^2+^-dependent neurotransmitter release, PTPσ loss may lead to abnormal cognitive or emotional behavioral states, owing to a dysfunction in excitatory neural circuit properties.

The presynaptic release phenotype we observed at two different synapse types is in agreement with findings indicating that LAR-RPTPs are key hubs for presynaptic functions (Takahashi & Craig, 2013, Um & Ko, 2013). Thus, an investigation of the roles of PTPσ is required to determine the mechanisms by which the complex interaction network of PTPσ organizes the properties of specific neural circuits. Our results strongly suggest that this activity has significant implications for the dynamics of specific neural circuits (**Fig 7**). Strikingly, specific deletion of PTPσ in CA1 neurons impaired anxiety-like behavior without altering sociability in mice, suggesting that presynaptic PTPσ mediates specific behaviors in mice.

Despite our results clearly linking presynaptic PTPσ functions in specific brain regions with specific behaviors, the causal relationships between presynaptic phenotypes and deficits in specific behaviors caused by presynaptic PTPσ deletion remain to be directly and fully determined (**Fig 7**). Anatomical effects of PTPσ KO are similarly in distal and proximal subicular neurons, which are molecularly heterogeneous links to parallel streams for disparate cognitive processing (Cembrowski *et al*, 2018). Thus, in-depth investigations are needed to determine details of the molecular mechanisms of PTPσ (e.g. alternative splicing-mediated specific synaptic adhesion pathways) and to concomitantly identify input/output-defined circuit elements responsible for behavioral deficits of presynaptic PTPσ KO observed in this study. These investigations may improve the understanding of the molecular logics of the responsible neural circuit dynamics and behavior. Anatomical and electrophysiological analyses of PTPσ cKO presented in the current study should be extended to better understand the synapse-organizing role of PTPσ as a fundamental building block of all types of synapses. In summary, our study demonstrated that PTPσ is a key factor in tuning presynaptic properties responsible for coordinating excitation-to-inhibition balance in neural circuits and certain behaviors, likely by organizing neurotransmitter release machinery at excitatory synapses.

## Materials and Methods

### Construction of expression vectors

#### 1. PTPσ rescue constructs

The AAV constructs for PTPσ protein rescue were generated using previously described L-313 lentiviral constructs as templates (Han *et al*, 2018). AAVs encoding human PTPσ WT and all PTPσ mutants (ΔD2 and C1157S) were amplified by PCR and subsequently subcloned into the *Xba*I and *BamH*I sites of the vector, a gift from Dr. Hailan Hu (Li *et al*, 2013). *<u>2</u>*<u>. Others.</u> The plasmids pAAV-hSyn-ΔCre-GFP and pAAV-hSyn-Cre-GFP were from Dr. Thomas C. Südhof (Stanford University, Palo Alto, CA, USA); FSW-ΔCre and FSW-Cre were from Dr. Pascal S. Kaeser (Harvard University, Cambridge, MA, USA); and pCAGG-VGLUT1-Venus was from Dr. Frank Polluex (Columbia University, New York, NY, USA).

### Antibodies

PTPδ-specific antibodies were generated in our laboratory by immunizing rabbits with keyhole limpet hemocyanin (KLH)-conjugated synthesized peptides specific to human PTPδ (PLKKSRGKFIKPWESPDEME). Rabbits were immunized three times with the KLH-conjugated peptides, and their sera were collected. PTPδ-specific antibodies (JK179, 1 mg/ml; RRID: AB_2810232) were affinity-purified using Sulfolink columns (Pierce) on which the same peptides had been immobilized. Commercially obtained antibodies included: mouse monoclonal anti-GAD67 (clone 1G10.2; Millipore; RRID: AB_2278725), guinea pig polyclonal anti-VGLUT1 (Millipore; RRID: AB_2301751), rabbit polyclonal anti-VGLUT1 (Synaptic Systems; RRID: AB_887880), rabbit polyclonal anti-GABA_A_Rγ2 (Synaptic Systems; RRID: AB_2263066), mouse monoclonal anti-PSD-95 (clone K28/43; Neuromab; RRID: AB_2292909), mouse monoclonal anti-PTPσ (clone 17G7.2; MediMabs; RRID: AB_1808357), mouse monoclonal anti-CASK (clone K56A/50; NeuroMab; RRID: AB_2068730), mouse monoclonal anti-HA (clone 16B12; BioLegend; RRID: AB_2565006); mouse monoclonal anti-Bassoon (clone SAP7F407; Stressgen; RRID: AB_2313990); rabbit polyclonal anti-Munc13-1 (Synaptic Systems; RRID: AB_887733), rabbit polyclonal anti-RIM-BP2 (Synaptic Systems; RRID: AB_2619739), rabbit polyclonal anti-RIM1/2 (Synaptic Systems; RRID: AB_887775), mouse monoclonal anti-ELKS (Sigma-Aldrich; RRID: AB_2100013), mouse monoclonal anti-Synaptophysin (clone SVP-38; Sigma-Aldrich; RRID: AB_477523), mouse monoclonal anti-MAP2 (clone AP-20; Sigma-Aldrich; RRID: AB_477171), rabbit polyclonal anti-MAP2 (Abcam; RRID: AB_776174), mouse monoclonal anti-β-actin (clone C4; Santa Cruz Biotechnology; RRID: AB_626632), mouse monoclonal GluN1 (clone 54.1; Millipore; RRID: AB_94946), rabbit polyclonal Cav2.1 (Synaptic Systems; RRID: AB_2619841), and mouse monoclonal anti-gephyrin (clone 3B11; Synaptic Systems; RRID: AB_887717). Rabbit polyclonal anti-liprin α2 (RRID:AB_2810258) and rabbit polyclonal anti-liprin-α3 (RRID:AB_2810259) antibodies were gifts of Dr. Susanne Schoch-McGovern (Bonn, Germany); rat polyclonal anti-PTPδ antibody (RRID:AB_2810260) was the gift of Dr. Fumio Nakamura (Yokohama, Japan); rabbit polyclonal anti-pan-Shank (1172; RRID:AB_2810261), and rabbit polyclonal anti-Homer1 antibodies (1133; RRID: AB_2810985) were the gifts of Dr. Eunjoon Kim (KAIST, Korea).

### Chemicals

6-Cyano-7-nitroquinoxaline-2,3-dione (CNQX) was obtained from Sigma-Aldrich (Cat No. C127). Tetrodotoxin (TTX; Cat No. 1069); picrotoxin (Cat No. 1128), QX-314 (Cat No. 1014); and D-2-amino-5-phosphonovalerate (D-AP5; Cat No. 0106) were purchased from Tocris.

### Neuron culture, transfections, imaging, and quantitation

Hippocampal and cortical mouse neuron cultures were prepared from embryonic day 17 (E1) mouse embryos, as described previously (Ko *et al*, 2011). Mouse cultured neurons were seeded onto coverslips coated with poly-D-lysine (Sigma-Aldrich), and grown in Neurobasal medium supplemented with B-27 (Thermo Fisher), 0.5% FBS (WELGENE), 0.5 mM GlutaMAX (Thermo Fisher), and sodium pyruvate (Thermo Fisher). Cultured neurons were infected with lentiviruses at DIV3–4. For immunocytochemistry, cultured neurons were fixed with 4% paraformaldehyde/4% sucrose in PBS for 10–30 min at 4°C, and permeabilized with 0.2% Triton X-100 in PBS for 10–30 min at 4°C. Neurons were blocked with 3% horse serum/0.1% BSA in PBS for 15 min at room temperature and incubated with primary and secondary antibodies in blocking solution for 70 min at room temperature. The primary antibodies were used in these experiments included anti-VGLUT1 (Synaptic Systems; 1:700), anti-GAD67 (Millipore; 1:100), anti-GABA_A_Rγ2 (Synaptic Systems; 1:500), anti-GluA1 (1193; 1:200), anti-Gephyrin (Synaptic Systems; 1:100), and anti-pan-Shank (1172; 1:200). Images of randomly selected neurons were acquired using a confocal microscope (LSM800, Carl Zeiss) with a 63 × objective lens; all image settings were kept constant during image acquisition. Z-stack images obtained by confocal microscopy were converted to maximal projections, and puncta size and the density of the indicated presynaptic marker proteins were analyzed in a blinded manner using MetaMorph software (Molecular Devices Corp.).

### Production of lentiviruses

Lentiviruses were produced by transfecting HEK293FT cells with three plasmids—lentivirus vectors, psPAX2 (or pCMVdeltaR8.2), and pMD2.G—at a 2:2:1 ratio. After 72 h, lentiviruses were harvested by collecting the media as previously described (Han *et al*, 2018, Hsia *et al*, 2014).

### Production of adeno-associated viruses

HEK293T cells were co-transfected with the indicated AAV vectors, pHelper and AAV1.0 (serotype 2/9) capsids vectors. After 72 hours, the transfected HEK293T cells were collected, and resuspended in PBS, and lysed by subjecting them to four freeze-thaw cycles in an ethanol/dry ice bath (7 minutes each) and a 37°C water bath (5 min). The lysates were centrifuged and the supernatants were mixed with 40% polyethylene glycol and 2.5 M NaCl and centrifuged at 2000 × g for 30 min. The cell pellets were resuspended in HEPES buffer (20 mM HEPES, 115 mM NaCl, 1.2 mM CaCl_2_, 1.2 mM MgCl_2_, and 2.4 mM KH_2_PO_4_, pH 8.0) to which was added an equal volume of chloroform. The mixture was centrifuged at 400 × g for 5 min and concentrated three times with a Centriprep centrifugal filter (Cat. 4310, Millipore) at 1,220 × g (20 min each) and an Amicon Ultra centrifugal filter (Cat. UFC500396, Millipore) at 16,000 × g for 30 min. AAVs were titered by treating 1 μl of concentrated, filter-sterilized AAVs with 1 μl of DNase I (AMPD1; Sigma) for 30 min at 37 °C to eliminate any contaminating plasmid DNA. After treatment with 1 μl of stop solution (50 mM ethylenediaminetetraacetic acid) for 10 min at 65 °C, 10 μg of protease K (Cat. P2308; Sigma) was added and the sample was incubated for 1 h at 50°C. Reactions were stopped by heat inactivation at 95 °C for 20 min. The final virus titer was quantified by qRT-PCR. Empty AAV vector was used to generate a standard curve for qRT-PCRs targeting *GFP* sequences.

### qRT-PCRs

Cultured rat cortical neurons were infected with recombinant lentiviruses at DIV4 and harvested at DIV11 for qRT-PCR using SYBR green qPCR master mix (TaKaRa). Total RNA was extracted from mouse cortical neurons using TRIzol reagent (Invitrogen) according to the manufacturer’s protocol. Briefly, cells in each well of a 12-well plate of cultured neurons were harvested and incubated with 500 μl TRIzol reagent at room temperature for 5 minutes. After phenol-chloroform extraction, RNA in the upper aqueous phase was precipitated. cDNA was synthesized from 500 ng of RNA by reverse transcription using a ReverTra Ace-α kit (Toyobo). Quantitative PCR was performed on a CFX96 Touch Real-Time PCR system BioRad) using 0.5 μl of cDNA. The ubiquitously expressed β-actin was used as an endogenous control. The sequences of the primer pairs used were: mouse *Ptprs*, 5’-ATCAGAGAGCCCAAGGATCA-3’ (forward) and 5’-GCCACACACTCGTACACGTT-3’ (reverse); mouse *Ptprd*, 5’-CTCCTTGATCCCCATCTCTG–3’ (forward) and 5’-CAG GGCAGCCACTAAACTTC-3’ (reverse); and mouse *Ptprf*, 5’-CCCGATGGCTGAGTACAACA-3’ (forward) and 5’-CATCCCGGGCGTCTGTGA-3’ (reverse).

### Quantification of cell body size and Sholl analysis

Cultured mouse neurons were infected with lentivirus expressing EGFP-ΔCre or EGFP-Cre at DIV4, and the neurons were transfected with L-317 vector expressing mCherry protein at DIV8. Fluorescent images were acquired at DIV 11 using by confocal microscopy (LSM800; Carl Zeiss). The sizes of green fluorescence positive neurons were measured using MetaMorph software (Molecular Devices), and branch intersections were measured using Sholl analysis function of Fiji/ImageJ software (National Institute of Health, USA).

### Electron microscopy

E17 embryonic hippocampi of PTPσ and PTPδ mice were seeded onto 35 mm coverslips at densities of 450,000 cells/well. The neurons were infected with lentiviral vectors expressing ΔCre or Cre at DIV4. At DIV14, cultured neurons were fixed in 2% glutaraldehyde, 0.1 M Na-cacodylate buffer, pH 7.4, for 1 h at room temperature and overnight at 4°C. The cells were post-fixed in 0.5% OsO_4_ (osmium tetroxide), 0.8% K ferricyanide at room temperature for 60 min. All specimens were stained *en bloc* with 2% aqueous uranyl acetate for 30 min, dehydrated in a graded ethanol series up to 100%, embedded in Embed 812 resin (Electron Microscopy Science, PA), and polymerized overnight in a 60 °C oven. Thin sections (50–60 nm) were cut with a Leica ultramicrotome and post-stained with uranyl acetate and lead citrate. Sample grids were examined using a FEI Tecnai BioTWIN transmission electron microscope running at accelerating voltage of 80 kV. Images were recorded with a Morada CCD camera and iTEM (Olympus) software. This protocol allowed the unambiguous staining of membranes of synaptic vesicles as well as of pre- and post-synaptic compartments, resulting in accurate measurements of the nanoscale organization of the synaptic vesicles within nerve endings. To analyze synapse ultrastructure, the lengths of active zone and PSD, tethered vesicles, the membrane proximal vesicles, and total vesicle numbers were quantified using MetaMorph software (Molecular Devices). The numbers of total vesicles and docked vesicles were counted manually, and the distances from the active zone and the PSD to the vesicle center were measured. Vesicles located below 200 nm were considered membrane-proximal vesicles.

### Animals

PTPσ conditional knockout mice were purchased from The KOMP Repository Collection (UC Davis, USA). PTPδ conditional knockout mice were generated at Biocytogen Co., Ltd (Beijing, China). Ai9 reporter mice were purchased from Jackson Research Laboratories (007909). Nestin-Cre (003771, Jackson Research Laboratories) mice were the gift of Dr. Albert Chen (DUKE-NUS, Singapore). Wfs1-Cre mice (009103, Jackson Research Laboratories) were the gift of Dr. Susumu Tonegawa (Massachusetts Institute of Technology, USA). All mice were maintained and handled in accordance with protocols (DGIST-IACUC-17122104-001) approved by the Institutional Animal Care and Use Committee of DGIST. Mice were maintained under standard, temperature-controlled laboratory conditions on a 12:12 light/dark cycle (lights on at 9:00 am and off at 9:00 pm), and with water and food supplied *ad libitum.* All animal experiments were performed according to approved animal protocols at DGIST Laboratory Animal Resource Center.

### RNAscope analyses

RNAscope analyses of mouse brains were performed using RNAscope® Fluorescent Multiplex Assay kits (Advanced Cell Diagnostics) according to the manufacturer’s direction. Briefly, within 5 min of dissection, mouse brains were immersed in cryo-embedding medium and frozen on dry ice. Brain tissue was sliced into 20 μm-thick coronal sections using a cryotome (Model CM-3050-S; Leica Biosystems), mounted, and dried at –20°C for 10 min. Tissue samples were fixed with 4% formaldehyde for 15 minutes at 4°C and dehydrated by incubation at room temperatures (RT) in 50% EtOH for 5 min, in 70% EtOH for 5 min, and twice 100% EtOH for 5 min. The fixed samples were treated with protease IV for 30 min at RT and washed twice with 1X PBS. To detect RNA, the sections were incubated in different amplifier solutions in a HybEZ hybridization oven (Advanced Cell Diagnostics) at 40°C. Three synthetic oligonucleotides complementary to nucleotide residues 1051–1947 of Mm–*Ptprs*–C1, 1329–2486 of Mm-*Ptprd*–C1 and Mm-*Ptprd*–C2, and 4001–5386 of Mm–*Ptprf*-C3 (Advanced Cell Diagnostics) were labeled by conjugation to Alexa Fluor 488, Altto 550 and Altto 647, and the labeled probe mixtures were hybridized by tissue samples by incubating them with slide-mounted sections for 2 hours at 40°C. Nonspecifically hybridized probes were removed by washing the sections three times for 2 minutes each with 1X wash buffer at RT, followed by incubations at 40°C with Amplifier 1-FL for 30 minutes, Amplifier 2-FL for 15 minutes, Amplifier 3-FL for 30 minutes, and Amplifier 4 Alt B-FL for 15 minutes. Each amplifier was removed by washing twice in 1X wash buffer at RT. The fluorescence images were acquired using a LSM 800 microscope (Carl Zeiss).

### Stereotaxic surgery and virus injections

For behavioral and electrophysiology experiments, 4–5-week-old mice were anesthetized by intraperitoneal injection of 2% 2,2,2-tribromoethanol (Sigma), dissolved in saline, and secured in a stereotaxic apparatus. For immunohistochemistry (IHC), 6 to 7-week-old mice were used. Viral solutions were injected using a Nanoliter 2010 Injector (World Precision Instruments), including a NanoFil syringe and 33 gauge needle, at a flow rate of 50 nl/min (injected volume, 500 nl). The coordinates used for stereotaxic injections targeting the ventral hippocampal CA1 were, relative to the bregma, anteroposterior (AP) −3.1 mm; medial–lateral (ML), ± 3.2 mm; and dorsal–ventral (DV), −2.5 mm. The coordinates for targeting the mPFC were, relative to the bregma, AP, +1.7 mm; ML, ± 0.4 mm; and DV, −2.2 mm. Behavioral tests were performed 5 weeks after each injected mouse was returned to its home cage, and immunohistochemical and electrophysiological analyses were performed 3 weeks later.

### Immunohistochemistry

Male mice aged 8–10-weeks were anesthetized and immediately perfused, first with PBS for 5 minutes and then with 4% paraformaldehyde for 5 minutes. Their brains were removed, fixed overnight in 4% paraformaldehyde, incubated overnight in 30% sucrose (in PBS), and sliced into 35-μm-thick coronal sections using a cryotome (Model CM-3050-S; Leica Biosystems). The sections were permeabilized in PBS containing 0.5% Triton X-100 for 1 h and blocked in PBS containing 5% bovine serum albumen and 5% horse serum for 1 minutes. The brain sections were incubated overnight with primary antibodies for overnight at 4 °C. The following primary antibodies were used: anti-VGLUT1 (1:200), anti-GAD67 (1:100). The brain sections were washed three times with PBS and incubated with the appropriate Cy3-conjugated secondary antibodies (Jackson ImmunoResearch) for 2 hours at RT. After three washes with PBS, the sections were counterstained with DAPI (4’,6-diamidino-2-phenylinodole) and mounted onto glass slides (Superfrost Plus; Fisher Scientific) with Vectashield mounting medium (H-1200; Vector Laboratories).

### *In vitro* and *ex vivo* electrophysiology

<u>*1.Electrophysiology of primary cultured neurons*</u>. Hippocampal neurons obtained from PTPσ and PTPδ cKO mice were infected on DIV4 with lentiviruses encoding Cre-EGFP or dCre-EGFP, followed by analysis at DIV13-16. Pipettes were pulled from borosilicate glass (o.d. 1.5mm, i.d. 0.86mm; Sutter Instrument), using a Model P-97 pipette puller (Sutter Instrument). The resistance of pipettes filled with internal solution varied between 3-6 MΩ. The internal solution contained 145 mM Cs-methanesulfonate, 5 mM NaCl, 5 mM TEA-Cl, 10 mM HEPES, 5 mM EGTA, 0.3 mM Na-GTP, 4 mM Mg-ATP, 10 mM disodium phosphocreatine and 1 mM QX-314 with pH adjusted to 7.2–7.4 with KOH, and an osmolarity of 290–295 mOsmol/L. The external solution consisted of 130 mM NaCl, 4 mM KCl, 2 mM CaCl_2_, 1 MgCl_2_, 10 mM HEPES, and 10 mM D-glucose with pH adjusted to 7.2–7.4 with NaOH, and an osmolarity of 300–305 mOsmol/L. Whole-cell configuration was generated at RT using MPC-200 manipulators (Sutter Instrument) and a Multiclamp 700B amplifier (Molecular Devices). mEPSCs, mIPSCs, and sucrose EPSCs were recorded at a holding potential of −70 mV. For sucrose puffing, 500 mM sucrose was applied directly on the dendritic field of the patched neurons at a puff pressure of 6–8 psi using a PV-820 Pneumatic Picopump system (World Precision Instruments). Receptor-mediated synaptic responses were pharmacologically isolated by applying drug combinations of 50 µM picrotoxin, 10 µM CNQX, 50 µM D-APV and/or 1 µM tetrodotoxin. Synaptic currents were analyzed offline using Clampfit 10.5 (Molecular Devices) software. <u>*2.Acute slice electrophysiology*</u>. Transverse hippocampal formation and coronal mPFC (300 μm) were prepared from 10– 12-week-old male mice, as described (Noh *et* al, 2019). The mice were anesthetized with isoflurane and decapitated, and their brains were rapidly removed and placed in ice-cold, oxygenated (95% O_2_/5% CO_2_), low-Ca^2+^/high-Mg^2+^ dissection buffer containing 5 mM KCl, 1.23 mM NaH_2_PO_4_, 26 mM NaHCO_3_, 10 mM dextrose, 0.5 mM CaCl_2_, 10 mM MgCl_2_, and 212.7 mM sucrose. Slices were transferred to a holding chamber in an incubator containing oxygenated (95% O_2_/5% CO_2_) artificial cerebrospinal fluid (ACSF) containing 124 mM NaCl, 5 mM KCl, 1.23 mM NaH_2_PO_4_, 2.5 mM CaCl_2_, 1.5 mM MgCl_2_, 26 mM NaHCO_3_, and 10 mM dextrose at 28–30°C for at least 1 h before recording. After > 1 h incubation in ACSF, slices were transferred to a recording chamber with continuous perfusion (2 ml/min) by ACSF oxygenated with 95% O_2_/5% CO_2_ at 23–25°C. All recordings were performed on pyramidal neurons in the subiculum, mPFC layer V, or hippocampal CA1 area identified by their size and morphology. Virus-infected neurons were identified by GFP fluorescence. Patch pipettes (4–6 MΩ) were filled with a solution containing 130 mM Cs-MeSO_4_, 0.5 mM EGTA, 5 mM TEA-Cl, 8 mM NaCl, 10 mM HEPES, 1 mM QX-314, 4 mM ATP-Hg, 0.4 mM GTP-Na, 10 mM phosphocreatine-Na_2_, 0.1 mM spermine (for measuring mEPSCs, EPSC-PPRs and IPSC-PPRs); 130 mM CsCl, 1.1 mM EGTA, 2 mM MgCl_2_, 0.1 mM CaCl_2_, 10 mM NaCl, 10 mM HEPES, 2 mM ATP-Na, pH 7.4 with an osmolarity of 280–290 mOsmol/L (for measuring mIPSCs). The extracellular recording solution consisted of ACSF supplemented with picrotoxin (100 μM), TTX (1 μM), and DL-AP5 (50 μM) for measuring mEPSCs; picrotoxin (100 μM) for measuring EPSC-PPRs; TTX (1 μM), CNQX (20 μM) and DL-AP5 (50 μM) for measuring mIPSCs; and CNQX (20 μM) and DL-AP5 (50 μM) for measuring IPSC-PPRs. Evoked synaptic responses were elicited by stimulation (0.2 ms current pulses) using a concentric bipolar electrode (for CA1-subiculum synapses) or theta glass capillaries filled with ACSF (for mPFC layer II–V synapses) placed 200–300 mm in front of postsynaptic pyramidal neurons at intensities that produced 40–50% of the maximal E/IPSC amplitude. Recordings were obtained using a Multiclamp 700B amplifier (Molecular Devices) under visual control with differential interference contrast illumination on an upright microscope (BX51WI; Olympus). Only cells with an access resistance < 20 MΩ and an input resistance >100 MΩ were studied. The cells were discarded if the input or the access resistance changed more than 20%. Data were acquired and analyzed using pClamp 10.7 (Molecular Devices). Signals were filtered at 3 kHz and digitized at 10 kHz with DigiData 1550 (Molecular Devices).

### Mouse behavioral tests

Male PTPσ^f/f^::Wfs1-Cre (*Wfs1*-PTPσ), PTPσ^f/f^ (Ctrl; 7–10 weeks old), and PTPσ^f/f^ (9–11 weeks old) mice injected with AAV-ΔCre/EGFP (Control) or AAV-Cre/EGFP (PTPσ-cKO) into their CA1 regions were used for behavioral tests. The order of testing was Laboras, Y-maze, open-field, novel object-recognition, elevated-plus maze, light-dark transition (LDT), contextual fear conditioning (CFC) and three-chamber tests. Mice were excluded from quantitative analyses if one or both of the injections were off-target, as demonstrated by *post hoc* immunostaining after behavioral analyses.

### Behavioral analyses

<u>*1. Y-maze test*</u>. Mice were introduced into the center of a Y-shaped white acrylic maze with three 40-cm–long arms at 120° angles from each other and allowed to explore freely for 8 minutes. Entry into an arm was defined as the entry of all four limbs of a mouse. Mouse movement was recorded using a top-view infrared camera, and analyzed using EthoVision XT 10 software (Noldus). <u>*2. Open-field test*</u>. Mice were placed into a white acrylic open-field box (40 × 40 × 35 cm), and allowed to freely explore the environment for 30 minutes in the dark (0 lux). The distance moved and time spent in the center zone were recorded with a top-view infrared camera, and analyzed using EthoVision XT 10 software (Noldus). <u>*3. Novel object-recognition test.*</u> Mice were habituated to an open field chamber for 10 minutes. During training sessions, two identical objects were placed in the center of the chamber at regular intervals, and mice were allowed to explore the objects for 10 minutes. The mice were subsequently returned to their home cages for 24 hours. For novel object-recognition tests, one of the two objects was replaced by a new object placed in the same position of the chamber. Mice were returned to the chamber and allowed to explore freely for 10 minutes. The movement of mice was recorded by an infrared camera, and the number and duration of contacts were analyzed using EthoVision XT 10 (Noldus). <u>*4. Elevated plus-maze test.*</u> The elevated plus-maze is a plus-shaped (+) white acrylic maze with two open arms (30 × 5 × 0.5 cm) and two closed arms (30 × 5 × 30 cm) positioned at a height of 45 cm from the floor. Light conditions around open and closed arms were ∼300 and ∼30 lux, respectively. For the test, mice were introduced into the center zone of the elevated plus-maze and allowed to move freely for 5 minutes. All behaviors were recorded with a top-view infrared camera, and the time spent in each arm and the number of arm entries were measured and analyzed using EthoVision XT 10 software (Noldus). <u>5. *Light-dark transition test.*</u> The light-dark transition test maze consists of equally sized light and dark chambers, measuring 20 × 10 × 35 cm each. The light conditions for the light and dark chambers were 500∼600 and 0 lux, respectively. A mouse was introduced into the center of the light chamber and allowed to explore freely for 10 min. All behaviors were recorded by a top-view infrared camera, and the time spent and number of entries into the light chamber were measured and analyzed using EthoVision XT 10 software (Noldus). <u>*6. Three-chamber test.*</u> Mice were placed into a white acrylic box divided into three chambers, measuring 20 × 40 × 22 cm each, and allowed to freely explore the environment for 10 minutes in the dark (0 lux). After a habituation time, an age-matched social target mouse (S1) and an object target (O) were placed in wire cups on each side of the chamber, and the mouse nose point approaches to the mouse and object were measured for 10 min. The object target was replaced by a new social target mouse (S2) and social novelty was again measured for 10 min. The approach time was calculated as the times spent sniffing S1 and O, or S2 and S. <u>*7. Contextual fear conditioning test.*</u> Contextual fear conditioning tests were performed using circadian cabinets (Actimetrics). On training day, mice were placed in a fear conditioning chamber and allowed to habituate for 120 sec. Subsequently a tone (85 dB, 20 sec) paired with a footshock (0.55mA, 2 sec) separated by 1 min intervals was delivered three times. The mice were returned to their home cages for 24 h and transferred to the same conditioning chamber. Freezing times were measured for 180 sec after habituation for 120 sec. Altered context tests were performed 24 h later. During these tests, the conditioning chamber was modified with a white circular plastic cylinder on a white background. Freezing times were measured for 180 sec after habituation for 120 sec. After 3 h, freezing times were measured under a conditioning noise background for 180 sec after habituation for 120 sec. All freezing times were analyzed by FreezeFrame Ver1 (Coulbourn Instruments and ActiMetrics Fear Conditioning System). *<u>8. Laboras test.</u>* Mice were placed in a Laboratory Animal Behavior Observation Registration and Analysis System (Laboras System), and home cage activity (locomotion, climbing, rearing, grooming, eating and drinking) was monitored continuously for 48 h with a 12/12 light (9:00 am to 9:00 pm) /dark (9:00 pm to 9:00 am) cycle and *ad libitum* feeding.

## Acknowledgements

We thank Drs. Susumu Tonegawa (MIT, USA) and Albert Chen (DUKE-NUS, Singapore) for the gift of Wfs1-Cre and Nestin-Cre driver lines, respectively. This study was supported by the Korea Healthcare Technology R & D Project, funded by the Ministry for Health and Welfare Affairs, Republic of Korea (Grant HI17C0080 to J.K.).

## Author contributions

S.Y.C. and J.K. conceived the project; K.A.H, H.Y.L., D.L., J.S., T.H.Y., C.L. and X.L. performed the experiments; K.A.H., H.Y.L., D.L., J.S., T.H.Y., C.L., J.S.R., J.W.U., S.Y.C. and J.K. analyzed the data; S.Y.C. and J.K. wrote the manuscript with input from the other authors.

## Expanded View Figure Legends

**Expanded View Figure 1. Generation of PTP cKO mice**

**A, B** Conditional KO (cKO) strategy for PTPσ (**A**) and PTPδ (**B**) mouse lines. Exon 4 of the PTPσ gene and exon 12 of the PTPδ gene were targeted (left). Primer locations for the WT and post Flp alleles are indicated with arrows (middle). PCR genotyping of WT and PTP floxed mice (right).

**C** Quantitative RT-PCR analysis of neuron RNA. Relative levels of PTPσ, PTPδ, and LAR mRNAs were measured in cultured cortical neurons infected with lentiviruses expressing Cre-recombinase. Data are means ± SEMs (n = 4 independent experiments).

**D** Representative immunoblot analysis of level of PTPσ and PTPδ proteins in brain homogenates from 8-week old control and PTP cKO mice. Levels of PTPσ and PTPδ proteins were measured in the indicated PTP floxed mice crossed with *Nestin-Cre* mice (Nestin-*PTPσ* or Nestin-*PTPδ*) or respective PTP floxed mice (Ctrl). Arrows indicate band(s) immunoreactive with PTPσ-specific and PTPδ-specific (JK179) antibodies.

**E** Images illustrating the body size of littermate control (Ctrl), Nestin-*PTPσ* and Nestin-*PTPδ* mice at 2 months of age. Nestin-*PTPσ* and Nestin-*PTPδ* mice were significantly smaller than age- and sex-matched Ctrl mice.

**F** Birth rates of genotyped pups from breeding of PTPσ^f/f^ or PTPδ^f/f^ mice with the corresponding Nestin-*PTPσ^f/+^* or Nestin-PTPδ *^f/+^*mice. Data are means ± SEMs (n = 79 for PTPσ and 81 for PTPδ mice).

**Expanded View Figure 2. Deletion of PTPσ and PTPδ from hippocampal CA1 specifically decreases innervation of excitatory and inhibitory synaptic inputs on subicular neurons, respectively**

**A, D** Representative VGLUT1 (**A**) and GAD67 (**D**) positive immunofluorescence images of proximal and distal SuB of *PTPσ^f/f^* mice injected with AAV-ΔCre or AAV-Cre. Scale bar: 20 μm. **B, E** Quantification of the density, size and integrated intensity of VGLUT1-positive (**B**) and GAD67-positive (**E**) synaptic puncta in proximal SuB. Data are means ± SEMs (n denotes the number of analyzed brain slices; 18–19 brain slices from 4 mice; \*\**p* < 0.01; Mann Whitney U-test).

**C, F** Quantification of the density, size and integrated intensity of VGLUT1-positive (**C**) and GAD67-positive (**F**) synaptic puncta in distal SuB. Data are means ± SEMs (n denotes the number of analyzed brain slices; 18–19 brain slices from 4 mice; Mann Whitney U-test).

**G, J** Representative VGLUT1 (**G**) and GAD67 (**J**) positive immunofluorescence images of proximal and distal SuB of *PTPδ^f/f^* mice injected with AAV-ΔCre or AAV-Cre. Scale bar: 20 μm.

**H, K** Quantification of the density, size and integrated intensity of VGLUT1-positive (**H**) and GAD67-positive (**K**) synaptic puncta in proximal SuB. Data are means ± SEMs (n denotes the number of analyzed brain slices; 13–14 brain slices from 3 mice; \**p* < 0.05; \*\**p* < 0.01; Mann Whitney U-test).

**I, L** Quantification of the density, size and integrated intensity of VGLUT1-positive (**I**) and GAD67-positive (**L**) synaptic puncta in distal SuB. Data are means ± SEMs (n denotes the number of analyzed brain slices; 13–14 brain slices from 3 mice; Mann Whitney U-test).

**Expanded View Figure 3. Marginal effect of presynaptic deletion of PTPσ on excitatory and inhibitory synaptic transmission in pyramidal neurons of mPFC layer V and hippocampal subiculum**

**A–C** Representative sEPSC traces (**A**) recorded from mPFC layer V pyramidal neurons in acute mPFC slices from littermate control and Wfs1-*PTPσ* mice, and cumulative distribution of sEPSC frequencies (**B**) and amplitudes (**C**). Insets show average sEPSC frequencies (**B**) and amplitudes (**C**). Data are means ± SEMs (n denotes the number of analyzed neurons; Control, 13; and Wfs1-*PTPσ*, 15; two-tailed Student’s t-tests).

**D–F** Representative mIPSC traces (**D**) recorded from mPFC layer V pyramidal neurons in acute mPFC slices from littermate control and Wfs1-*PTPσ* mice, and cumulative distribution plots of mIPSC frequencies (**E**) and amplitudes (**F**). Insets show average mIPSC frequencies (**E**) and amplitudes (**F**). Data are means ± SEMs (n denotes the number of analyzed neurons; Control, 14; and Wfs1-*PTPσ*, 12; two-tailed Student’s t-tests).

**G–I** Representative sEPSC traces (**G**) recorded from SuB pyramidal neurons in acute SuB slices from littermate control and Wfs1-*PTPσ* mice, and cumulative distribution of sEPSC frequencies

(**H**) and amplitudes (**I**). Insets show average sEPSC frequencies (**H**) and amplitudes (**I**). Data are means ± SEMs (n denotes the number of analyzed neurons; Control, 12; and Wfs1-*PTPσ*, 21; two-tailed Student’s t-tests).

**J–L** Representative mEPSC traces (**J**) recorded from CA1 pyramidal neurons in acute CA1 slices from littermate control and Wfs1-*PTPσ* mice, and cumulative distribution plots of mEPSC frequencies (**K**) and amplitudes (**L**). Insets show average mEPSC frequencies (**K**) and amplitudes (**L**). Data are means ± SEMs (n denotes the number of analyzed neurons; Control, 11; and Wfs1-*PTPσ*, 13; two-tailed Student’s t-tests).

**Expanded View Figure 4. Analysis of PTPσ extracellular and intracellular mechanisms involved in controlling innervation of excitatory inputs in subiculum**

**A** Schematic diagram of AAV constructs expressing PTPσ WT and the indicated PTPσ variants and experimental strategy (see Han *et al*, 2018).

**B** Representative images of subiculum 3 weeks after stereotactic infection of Wfs1-*PTPσ* mice with the indicated AAVs, followed by immunostaining for EGFP (green) to detect infected neurons and HA to detect the expression of AAV-rescue viruses (red). Scale bar, 20 μm.

**C** Representative immunofluorescence images of the proximal and distal subiculum of Wfs1-*PTPσ* mice 3 weeks after stereotactic injection of the indicated AAVs. Brain sections were immunostained for VGLUT1 and HA to show neurons infected with the indicated AAVs. Scale bar: 20 μm.

**D, E** Quantification of the density, size and integrated intensity of VGLUT1-positive synaptic puncta in proximal (**D**) and distal (**E**) subicular neurons. Data are means ± SEMs (n denotes the number of analyzed brain slices; n = 14–18 slices from 3 mice; \**p* < 0.05, ANOVA with a non-parametric Kruskal-Wallis test).

**Expanded View Figure 5. Wfs1-*PTPσ* mice exhibit no altered behaviors, including changes in locomotion, light-induced anxiety-like behavior, spatial working memory, learning and memory**

**A, B** Activities of control and Wfs1-*PTPσ* mice in Laboras cages, where movements of mice were monitored for 48 hours. Data are means ± SEMs (Control, n = 16; Wfs1-*PTPσ*, n =16; Mann-Whitney U Test).

**C, D** Analysis of the locomotor activity of Wfs1-*PTPσ* mice by open field tests. Representative heat map (**C**) and graphs (**D**) showing comparable locomotor activity of control and Wfs1-*PTPσ* mice, as recorded by distance moved, time spent in the center region of the open field arena, and frequency of entry into this region. Data are means ± SEMs (Control, n = 19; Wfs1-*PTPσ,* n = 15; Mann-Whitney U test).

**E, F** Analysis of the light-induced anxiety-like behavior by light-dark transition tests in Wfs1-*PTPσ* mice. Representative heat map (**E**) and graphs (**F**) showing comparable light-induced anxiety-like behavior of control and Wfs1-*PTPσ* mice, as measured by time spent in light box and number of total entries into the light chamber. Data are means ± SEMs (Control, n = 18; Wfs1-*PTPσ,* n = 18; Mann-Whitney U test).

**G, H** Analysis of the spatial working memory by Y-maze tests in Wfs1-*PTPσ* mice. Representative heat map (**E**) and graphs (**F**) showing comparable spatial working memory of control and Wfs1-*PTPσ* mice, as measured by spontaneous alternation performance ratio and total number of arm entries. Data are means ± SEMs (Control, n = 11; Wfs1-*PTPσ,* n = 11; Mann-Whitney U test).

**I, J** Analysis of object recognition memory by novel object recognition tests in Wfs1-*PTPσ* mice. Representative heat map (**I**) and graphs (**J**) showing comparable object recognition memory of control and Wfs1-*PTPσ* mice, as measured by exploration times during the training and testing phases. Data are means ± SEMs (Control, n = 11; Wfs1-*PTPσ,* n = 11; \*\**p* < 0.01; Mann-Whitney U test).

**K, L** Analysis of contextual fear memory by contextual fear conditioning tests in Wfs1-*PTPσ* mice. Design of the contextual fear conditioning test (**K**) and graphs (**L**) showing comparable contextual memory of control and Wfs1-*PTPσ* mice. Data are means ± SEMs (Control, n = 16; Wfs1-*PTPσ,* n = 16, \*\**p* < 0.01; Mann-Whitney U test).

## References

Abbott LF, Regehr WG (2004) Synaptic computation. Nature 431: 796–803

Ackley BD, Harrington RJ, Hudson ML, Williams L, Kenyon CJ, Chisholm AD, Jin Y (2005) The two isoforms of the Caenorhabditis elegans leukocyte-common antigen relate d receptor tyrosine phosphatase PTP-3 function independently in axon guidance and syna pse formation. J Neurosci 25: 7517–7528

Acuna C, Liu X, Südhof TC (2016) How to Make an Active Zone: Unexpected Univers al Functional Redundancy between RIMs and RIM-BPs. Neuron 91: 792–807

Aoto J, Foldy C, Ilcus SM, Tabuchi K, Südhof TC (2015) Distinct circuit-dependent fun ctions of presynaptic neurexin-3 at GABAergic and glutamatergic synapses. Nat Neurosci 18: 997–1007

Aoto J, Martinelli DC, Malenka RC, Tabuchi K, Südhof TC (2013) Presynaptic neurexin -3 alternative splicing trans-synaptically controls postsynaptic AMPA receptor trafficking. Cell 154: 75–88

Cembrowski MS, Phillips MG, DiLisio SF, Shields BC, Winnubst J, Chandrashekar J, B as E, Spruston N (2018) Dissociable Structural and Functional Hippocampal Outputs via Distinct Subiculum Cell Classes. Cell 173: 1280–1292

Chagnon MJ, Uetani N, Tremblay ML (2004) Functional significance of the LAR recept or protein tyrosine phosphatase family in development and diseases. Biochem Cell Biol 8 2: 664–675

Chanda S, Hale WD, Zhang B, Wernig M, Südhof TC (2017) Unique versus Redundant Functions of Neuroligin Genes in Shaping Excitatory and Inhibitory Synapse Properties. J Neurosci 37: 6816–6836

Chen LY, Jiang M, Zhang B, Gokce O, Südhof TC (2017) Conditional Deletion of All Neurexins Defines Diversity of Essential Synaptic Organizer Functions for Neurexins. Ne uron 94: 611–625

Choi Y, Nam J, Whitcomb DJ, Song YS, Kim D, Jeon S, Um JW, Lee SG, Woo J, K won SK, Li Y, Mah W, Kim HM, Ko J, Cho K, Kim E (2016) SALM5 trans-synaptica lly interacts with LAR-RPTPs in a splicing-dependent manner to regulate synapse develo pment. Sci Rep 6: 26676

Dai J, Aoto J, Südhof TC (2019) Alternative Splicing of Presynaptic Neurexins Different ially Controls Postsynaptic NMDA and AMPA Receptor Responses. Neuron 102: 993–100 8

Dai Y, Taru H, Deken SL, Grill B, Ackley B, Nonet ML, Jin Y (2006) SYD-2 Liprin-a lpha organizes presynaptic active zone formation through ELKS. Nat Neurosci 9: 1479–1 487

Doussau F, Augustine GJ (2000) The actin cytoskeleton and neurotransmitter release: an overview. Biochimie 82: 353–363

Dunah AW, Hueske E, Wyszynski M, Hoogenraad CC, Jaworski J, Pak DT, Simonetta A, Liu G, Sheng M (2005) LAR receptor protein tyrosine phosphatases in the developm ent and maintenance of excitatory synapses. Nat Neurosci 8: 458–467

Elchebly M, Wagner J, Kennedy TE, Lanctot C, Michaliszyn E, Itie A, Drouin J, Trem blay ML (1999) Neuroendocrine dysplasia in mice lacking protein tyrosine phosphatase s igma. Nat Genet 21: 330–333

Etherton MR, Blaiss CA, Powell CM, Südhof TC (2009) Mouse neurexin-1alpha deletio n causes correlated electrophysiological and behavioral changes consistent with cognitive impairments. Proc Natl Acad Sci U S A 106: 17998–18003

Grauel MK, Maglione M, Reddy-Alla S, Willmes CG, Brockmann MM, Trimbuch T, Ro senmund T, Pangalos M, Vardar G, Stumpf A, Walter AM, Rost BR, Eickholt BJ, Hauc ke V, Schmitz D, Sigrist SJ, Rosenmund C (2016) RIM-binding protein 2 regulates relea se probability by fine-tuning calcium channel localization at murine hippocampal synapse s. Proc Natl Acad Sci U S A 113: 11615–11620

Han KA, Jeon S, Um JW, Ko J (2016) Emergent Synapse Organizers: LAR-RPTPs and Their Companions. Int Rev Cell Mol Biol 324: 39–65

Han KA, Ko JS, Pramanik G, Kim JY, Tabuchi K, Um JW, Ko J (2018) PTPsigma Dri ves Excitatory Presynaptic Assembly via Various Extracellular and Intracellular Mechanis ms. J Neurosci 38: 6700–6721

Han KA, Um JW, Ko J (2019) Intracellular protein complexes involved in synapse asse mbly in presynaptic neurons. Adv Protein Chem Struct Biol 116: 347–373

Horn KE, Xu B, Gobert D, Hamam BN, Thompson KM, Wu CL, Bouchard JF, Uetani N, Racine RJ, Tremblay ML, Ruthazer ES, Chapman CA, Kennedy TE (2012) Receptor protein tyrosine phosphatase sigma regulates synapse structure, function and plasticity. J Neurochem 122: 147–161

Hsia HE, Kumar R, Luca R, Takeda M, Courchet J, Nakashima J, Wu S, Goebbels S, An W, Eickholt BJ, Polleux F, Rotin D, Wu H, Rossner MJ, Bagni C, Rhee JS, Brose N, Kawabe H (2014) Ubiquitin E3 ligase Nedd4-1 acts as a downstream target of PI3K/ PTEN-mTORC1 signaling to promote neurite growth. Proc Natl Acad Sci U S A 111: 1 3205–3210

Kaeser PS, Deng L, Wang Y, Dulubova I, Liu X, Rizo J, Südhof TC (2011) RIM prote ins tether Ca^2+^ channels to presynaptic active zones via a direct PDZ-domain interaction. Cell 144: 282–295

Kaeser PS, Kwon HB, Chiu CQ, Deng L, Castillo PE, Südhof TC (2008) RIM1alpha a nd RIM1beta are synthesized from distinct promoters of the RIM1 gene to mediate diffe rential but overlapping synaptic functions. J Neurosci 28: 13435–13447

Kitamura T, Pignatelli M, Suh J, Kohara K, Yoshiki A, Abe K, Tonegawa S (2014) Isla nd cells control temporal association memory. Science 343: 896–901

Ko J, Soler-Llavina GJ, Fuccillo MV, Malenka RC, Südhof TC (2011) Neuroligins/LRRT Ms prevent activity- and Ca^2+^/calmodulin-dependent synapse elimination in cultured neuro ns. J Cell Biol 194: 323–334

Ko JS, Pramanik G, Um JW, Shim JS, Lee D, Kim KH, Chung GY, Condomitti G, Ki m HM, Kim H, de Wit J, Park KS, Tabuchi K, Ko J (2015) PTPsigma functions as a presynaptic receptor for the glypican-4/LRRTM4 complex and is essential for excitatory synaptic transmission. Proc Natl Acad Sci U S A 112: 1874–1879

Kwon SK, Woo J, Kim SY, Kim H, Kim E (2010) Trans-synaptic adhesions between ne trin-G ligand-3 (NGL-3) and receptor tyrosine phosphatases LAR, protein-tyrosine phosph atase delta (PTPdelta), and PTPsigma via specific domains regulate excitatory synapse fo rmation. J Biol Chem 285: 13966–13978

Li K, Zhou T, Liao L, Yang Z, Wong C, Henn F, Malinow R, Yates JR, 3rd, Hu H (2013) betaCaMKII in lateral habenula mediates core symptoms of depression. Science 341 : 1016–1020

Li Y, Zhang P, Choi TY, Park SK, Park H, Lee EJ, Lee D, Roh JD, Mah W, Kim R, Kim Y, Kwon H, Bae YC, Choi SY, Craig AM, Kim E (2015) Splicing-Dependent Tran s-synaptic SALM3-LAR-RPTP Interactions Regulate Excitatory Synapse Development and Locomotion. Cell Rep 12: 1618–1630

Luuk H, Koks S, Plaas M, Hannibal J, Rehfeld JF, Vasar E (2008) Distribution of Wfs 1 protein in the central nervous system of the mouse and its relation to clinical sympto ms of the Wolfram syndrome. J Comp Neurol 509: 642–660

Madisen L, Zwingman TA, Sunkin SM, Oh SW, Zariwala HA, Gu H, Ng LL, Palmiter RD, Hawrylycz MJ, Jones AR, Lein ES, Zeng H (2010) A robust and high-throughput Cre reporting and characterization system for the whole mouse brain. Nat Neurosci 13: 133–140

McLean J, Batt J, Doering LC, Rotin D, Bain JR (2002) Enhanced rate of nerve regene ration and directional errors after sciatic nerve injury in receptor protein tyrosine phosph atase sigma knock-out mice. J Neurosci 22: 5481–5491

Missler M, Südhof TC, Biederer T (2012) Synaptic cell adhesion. Cold Spring Harb Pe rspect Biol 4: a005694

Mongillo G, Barak O, Tsodyks M (2008) Synaptic theory of working memory. Science 3 19: 1543–1546

Muller CS, Haupt A, Bildl W, Schindler J, Knaus HG, Meissner M, Rammner B, Striess nig J, Flockerzi V, Fakler B, Schulte U (2010) Quantitative proteomics of the Cav2 cha nnel nano-environments in the mammalian brain. Proc Natl Acad Sci U S A 107: 14950–14957

Nakamura F, Okada T, Shishikura M, Uetani N, Taniguchi M, Yagi T, Iwakura Y, Ohshi ma T, Goshima Y, Strittmatter SM (2017) Protein Tyrosine Phosphatase delta Mediates t he Sema3A-Induced Cortical Basal Dendritic Arborization through the Activation of Fyn Tyrosine Kinase. J Neurosci 37: 7125–7139

Neher E, Sakaba T (2008) Multiple roles of calcium ions in the regulation of neurotrans mitter release. Neuron 59: 861–872

Nelson JC, Stavoe AK, Colon-Ramos DA (2013) The actin cytoskeleton in presynaptic a ssembly. Cell Adh Migr 7: 379–387

Noh K, Lee H, Choi TY, Joo Y, Kim SJ, Kim H, Kim JY, Jahng JW, Lee S, Choi SY, Lee SJ (2019) Negr1 controls adult hippocampal neurogenesis and affective behaviors. Mol Psychiatry 24: 1189–1205

O’mara S (2005) The subiculum: what it does, what it might do, and what neuroanatomy has yet to tell us. J Anat 207: 271–282

Rosenmund C, Stevens CF (1996) Definition of the readily releasable pool of vesicles at hippocampal synapses. Neuron 16: 1197–1207

Schroeder A, Vanderlinden J, Vints K, Ribeiro LF, Vennekens KM, Gounko NV, Wierda KD, de Wit J (2018) A Modular Organization of LRR Protein-Mediated Synaptic Adhe sion Defines Synapse Identity. Neuron 99: 329–344

Spangler SA, Schmitz SK, Kevenaar JT, de Graaff E, de Wit H, Demmers J, Toonen R F, Hoogenraad CC (2013) Liprin-alpha2 promotes the presynaptic recruitment and turnov er of RIM1/CASK to facilitate synaptic transmission. J Cell Biol 201: 915–28

Südhof TC (2012) The presynaptic active zone. Neuron 75: 11–25

Südhof TC (2017) Synaptic Neurexin Complexes: A Molecular Code for the Logic of N eural Circuits. Cell 171: 745–769

Südhof TC (2018) Towards an Understanding of Synapse Formation. Neuron 100: 276–2 93

Takahashi H, Arstikaitis P, Prasad T, Bartlett TE, Wang YT, Murphy TH, Craig AM (2011) Postsynaptic TrkC and presynaptic PTPsigma function as a bidirectional excitatory sy naptic organizing complex. Neuron 69: 287–303

Takahashi H, Craig AM (2013) Protein tyrosine phosphatases PTPdelta, PTPsigma, and LAR: presynaptic hubs for synapse organization. Trends Neurosci 36: 522–534

Takahashi H, Katayama K, Sohya K, Miyamoto H, Prasad T, Matsumoto Y, Ota M, Yas uda H, Tsumoto T, Aruga J, Craig AM (2012) Selective control of inhibitory synapse d evelopment by Slitrk3-PTPdelta trans-synaptic interaction. Nat Neurosci 15: 389–98

Thompson KM, Uetani N, Manitt C, Elchebly M, Tremblay ML, Kennedy TE (2003) R eceptor protein tyrosine phosphatase sigma inhibits axonal regeneration and the rate of a xon extension. Mol Cell Neurosci 23: 681–92

Uetani N, Chagnon MJ, Kennedy TE, Iwakura Y, Tremblay ML (2006) Mammalian mot oneuron axon targeting requires receptor protein tyrosine phosphatases sigma and delta. J Neurosci 26: 5872–80

Uetani N, Kato K, Ogura H, Mizuno K, Kawano K, Mikoshiba K, Yakura H, Asano M, Iwakura Y (2000) Impaired learning with enhanced hippocampal long-term potentiation in PTPdelta-deficient mice. EMBO J 19: 2775–85

Um JW, Ko J (2013) LAR-RPTPs: synaptic adhesion molecules that shape synapse deve lopment. Trends Cell Biol 23: 465–75

Valnegri P, Montrasio C, Brambilla D, Ko J, Passafaro M, Sala C (2011) The X-linked intellectual disability protein IL1RAPL1 regulates excitatory synapse formation by bindin g PTPdelta and RhoGAP2. Hum Mol Genet 20: 4797–809

Wallace MJ, Batt J, Fladd CA, Henderson JT, Skarnes W, Rotin D (1999) Neuronal def ects and posterior pituitary hypoplasia in mice lacking the receptor tyrosine phosphatase PTPsigma. Nat Genet 21: 334–8

Wang SSH, Held RG, Wong MY, Liu C, Karakhanyan A, Kaeser PS (2016) Fusion Co mpetent Synaptic Vesicles Persist upon Active Zone Disruption and Loss of Vesicle Doc king. Neuron 91: 777–791

Wong MY, Liu C, Wang SSH, Roquas ACF, Fowler SC, Kaeser PS (2018) Liprin-alpha 3 controls vesicle docking and exocytosis at the active zone of hippocampal synapses. P roc Natl Acad Sci U S A 115: 2234–2239

Wyszynski M, Kim E, Dunah AW, Passafaro M, Valtschanoff JG, Serra-Pages C, Streuli M, Weinberg RJ, Sheng M (2002) Interaction between GRIP and liprin-alpha/SYD2 is r equired for AMPA receptor targeting. Neuron 34: 39–52

Yim YS, Kwon Y, Nam J, Yoon HI, Lee K, Kim DG, Kim E, Kim CH, Ko J (2013) Slitrks control excitatory and inhibitory synapse formation with LAR receptor protein tyr osine phosphatases. Proc Natl Acad Sci U S A 110: 4057–4062

Yoshida T, Yasumura M, Uemura T, Lee SJ, Ra M, Taguchi R, Iwakura Y, Mishina M (2011) IL-1 receptor accessory protein-like 1 associated with mental retardation and autis m mediates synapse formation by trans-synaptic interaction with protein tyrosine phospha tase delta. J Neurosci 31: 13485–99

Zhen M, Jin Y (1999) The liprin protein SYD-2 regulates the differentiation of presynaptic termini in C. elegans. Nature 401: 371–5

Zucker RS, Regehr WG (2002) Short-term synaptic plasticity. Annu Rev Physiol 64: 355–405

